# Outcome of H5N1 clade 2.3.4.4b virus infection in calves and lactating cows

**DOI:** 10.1101/2024.08.09.607272

**Authors:** Nico Joel Halwe, Konner Cool, Angele Breithaupt, Jacob Schön, Jessie D. Trujillo, Mohammed Nooruzzaman, Taeyong Kwon, Ann Kathrin Ahrens, Tobias Britzke, Chester D. McDowell, Ronja Piesche, Gagandeep Singh, Vinicius Pinho dos Reis, Sujan Kafle, Anne Pohlmann, Natasha N. Gaudreault, Björn Corleis, Franco Matias Ferreyra, Mariano Carossino, Udeni B.R. Balasuriya, Lisa Hensley, Igor Morozov, Lina M. Covaleda, Diego Diel, Lorenz Ulrich, Donata Hoffmann, Martin Beer, Juergen A. Richt

## Abstract

In March 2024, highly pathogenic avian influenza virus (HPAIV) clade 2.3.4.4b H5N1 infections in dairy cows were first reported from Texas, USA. Rapid dissemination to more than 190 farms in 13 states followed. Here, we provide results of two independent clade 2.3.4.4b experimental infection studies evaluating (i) oronasal susceptibility and transmission in calves to a US H5N1 bovine isolate genotype B3.13 (H5N1 B3.13) and (ii) susceptibility of lactating cows following direct mammary gland inoculation of either H5N1 B3.13 or a current EU H5N1 wild bird isolate genotype euDG (H5N1 euDG). Inoculation of the calves resulted in moderate nasal replication and shedding with no severe clinical signs or transmission to sentinel calves. In dairy cows, infection resulted in no nasal shedding, but severe acute mammary gland infection with necrotizing mastitis and high fever was observed for both H5N1 genotypes/strains. Milk production was rapidly and drastically reduced and the physical condition of the cows was severely compromised. Virus titers in milk rapidly peaked at 10^8^ TCID_50_/mL, but systemic infection did not ensue. Notably, adaptive mutation PB2 E627K emerged after intramammary replication of H5N1 euDG. Our data suggest that in addition to H5N1 B3.13, other HPAIV H5N1 strains have the potential to replicate in the udder of cows and that milk and milking procedures, rather than respiratory spread, are likely the primary routes of H5N1 transmission between cattle.

## Main

Epidemic occurrence of highly pathogenic avian influenza (HPAIV) of subtype H5 has developed, since 2022, into a panzootic with dynamic spread into an expansive number of host species^1–7^. In 2021, A/H5N1 clade 2.3.4.4b crossed the Atlantic and rapidly diffused through wild bird and commercial poultry populations in the Americas^3,8,9^. Subsequent reports of sporadic mammalian infections have become more frequent with data suggestive of mammal-to-mammal transmission chains present in South American seals since 2023^10,11^.

Historically, natural infections of cattle with influenza A virus (IAV) are not well documented^12^ despite the rare detection of IAV seropositive cattle^13^; but in March 2024, an outbreak of HPAIV H5N1 was reported in dairy cows in Texas caused by the novel B3.13 genotype, a reassortant of an ancestral European 2.3.4.4b virus and North American wild bird AIVs (H5N1 B3.13)^9^. Phylogenetic analyses of whole genome sequences recovered from wild birds, poultry, and mammals suggest a single spillover event into cattle, with the time to the most recent common ancestor indicating introduction occurring in late 2023 or early 2024^14,15^. Current epidemiological data suggests that subsequent inter-farm spread is mainly associated with unknowingly transporting infected cows^9^. As of 8^th^ August 2024, 190 dairy cattle farms in 13 US states have been affected^16^.

In the field, high level H5N1 B3.13 replication has been reported in the mammary gland of infected cows, resulting in high-titer virus shedding in milk, accompanied by mastitis, a massive drop in milk production, and limited reports of respiratory disease^9,17^. The susceptibility and rapid viral replication of HPAIV in the mammary gland are consistent with the evidence of highly abundant α2,3 linked sialic acids receptors in the bovine udder^18^. A novel PB2 M631L substitution accompanied the switch from avian to bovine hosts as a marker mutation^9,14^.

Spillover of bovine-origin B3.13 into several mammalian hosts (racoons, cats, etc.) has been reported^9,19^, as well as spillback into domestic and wild avian species with maintenance of bovine adaptations has been sporadically observed^14^. Recent human cases of H5N1 have also been directly linked to workers after having contact with affected cattle or poultry farms, causing conjunctivitis and conjunctival hemorrhage^20^. Accordingly, the current series of outbreaks in US cattle presents several urgent and unanswered questions: (i) Is the B3.13 genotype able to replicate in the bovine respiratory tract with viral shedding capable of onward transmission? (ii) At what timepoint after infection do cattle produce IAV-specific neutralizing antibodies? (iii) Is the mammary gland also permissive for infection with other H5N1 clade 2.3.4.4b strains? (iv) What is the clinical presentation, and what is the duration of virus shedding in milk? Finally, (v) does an H5N1 infection of the mammary gland lead to systemic spread?

Here, we performed two independent *in vivo* experiments to investigate the clinical outcome, pathogenicity, transmission, and tissue tropism of H5N1 clade 2.3.4.4b in calves and multiparous lactating cows. Calves (n=6) were oronasally inoculated with H5N1 B3.13 ^9^ and co-housed with sentinel animals (n=3), with additional calves serving as negative controls (n=3). The same virus isolate was used for an intramammary inoculation of lactating cows (n=3). For comparison, three additional lactating cows were inoculated with an EU genotype euDG H5N1 clade 2.3.4.4b wild bird virus isolate (H5N1 euDG). One lactating cow served as a negative control.

## Results

### Subclinical disease in calves oronasally infected with H5N1

Twelve healthy Holstein calves were enrolled in this study and allocated into three experimental groups: Principal-infected animals (n=6); sentinel animals (n=3); negative controls (n=3). Six principal-infected calves were oronasally inoculated with 1×10^6^ TCID_50_/calf of a virus suspension of H5N1 B3.13 (A/Cattle/Texas/063224-24-1/2024, GISAID accession number: EPI_ISL_19155861). Two days post infection, sentinel calves were co-mingled with principal-infected calves (Fig. 1A). All calves were monitored daily for clinical signs and clinical samples were collected at regular time-points (Fig 1A).

**Fig. 1.**
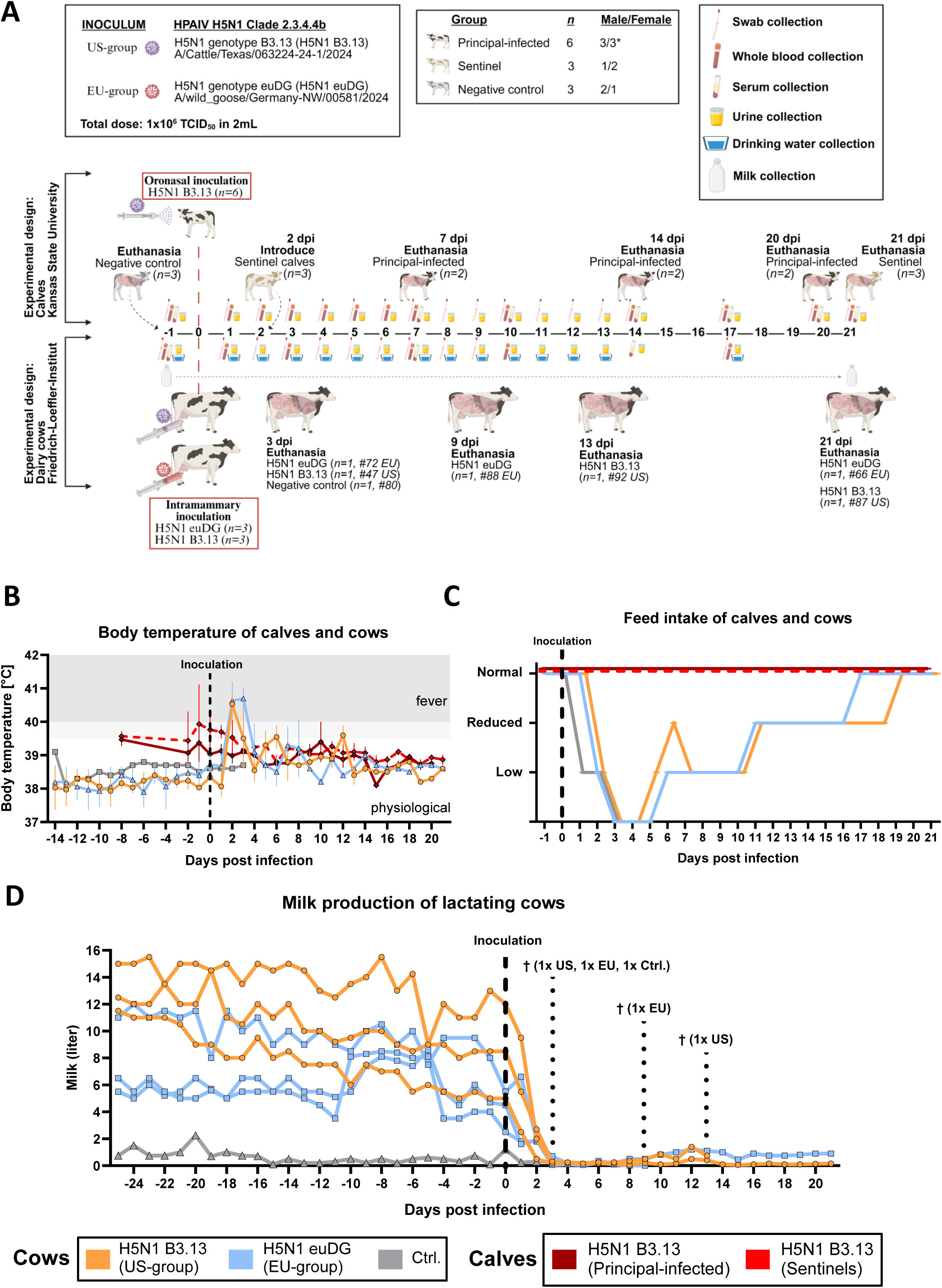
| Experimental design and clinical outcomes following infection with HPAIV H5N1 clade 2.3.4.4b isolates. **A** Experimental study timeline. (**Top**) Twelve Holstein calves of mixed sex (*\* indicates one calf was hermaphroditic*) were allocated to three experimental groups: 1 – principal-infected (n=6); 2 – sentinel (n=3); 3 – negative control (n=3). Negative control calves were euthanized prior to experimental infection and tissues were collected for baseline comparison. Principal-infected calves were oronasally inoculated with 1×10^6^ TCID_50_/calf of H5N1 B3.13. Sentinel calves were introduced 48 hours post-infection. Rectal temperatures and clinical samples, including whole blood, urine, nasal-, oral-, and rectal swabs, were collected daily for 14 dpi and every 3 days thereafter. Serum was collected at 0, 7, 10, 14, 17, and 20/21 dpi. Post-mortem examinations and extensive tissue collections were performed on days 7 (n=2, principal-infected), 14 (n=2, principal-infected), and 20/21 (n=2/3, principal-infected/sentinel) post infection. (**Bottom**) Seven Holstein-Friesian multiparous lactating dairy cattle were used in this experiment. Three animals were inoculated intramammary with 10^6.1^ TCID_50_/cattle of H5N1 B3.13 (A/Cattle/Texas/063224-24-1/2024, US-group, n=3) and three animals were inoculated intramammary with 10^5.9^ TCID_50_/cattle of H5N1 euDG (A/wild_goose/Germany-NW/00581/2024, EU-group, n=3). One cow served as a negative control. Swab samples (nasal, conjunctival, and rectal) were taken daily until 9 dpi. EDTA blood samples were taken from individual cattle at 1, 3, 7 and 10 dpi. Urine was taken regularly until 14 dpi. Serum samples were obtained from 7, 14 dpi and the day of euthanasia. One cow of each group (#47 US and #72 EU) reached the humane endpoint at 3 dpi, one further cow at 9 dpi (#88 EU) and one additional cow at 13 dpi (#92 US) and were subsequently subjected to necropsy. Created with BioRender under agreement number YH275PUF4T. **B** The mean and standard deviation of rectal temperatures are shown for both principal-infected and sentinel calves and each group of lactating cattle prior to and following inoculation. **C** The average feed intake of each group of calves and cows following H5N1-infection **D** Individual milk production of lactating cows prior to and following experimental infection. Milk production of individual cows was tracked daily from –25 dpi through until the end of the experiment at 21 dpi. The control animal never produced high amounts of milk, which is why this cow was picked as control. Dark red: Principal-infected calves (n=6). Bright red: Sentinel calves (n=3). Orange: H5N1 B3.13 infected lactating cows (US-group, n=3, #47, #87, #92) Blue: H5N1 euDG infected lactating cows (EU-group, n=3, #66, #72, #88) Grey: Uninfected negative control cow (#80).

Throughout the 21-day study period, signs of mild respiratory illness were occasionally observed in calves, including nasal mucus secretions (#712 at 2 days post infection (dpi); #754 at 8 and 9 dpi; #6772 at 2 and 6 dpi) and coughing (#6772 at 2 dpi; #754, persistent from 2 dpi until euthanasia). Rectal temperatures generally remained within normal range (Fig. 1B), and no other clinical signs consistent with acute illness, or consistent with clinical signs reported in impacted dairy cattle in the US were observed. All calves maintained normal appetite (feed intake) and normal activity levels (Fig. 1C).

### Severe disease in dairy cows caused by intramammary infection with two distinct H5N1 clade 2.3.4.4b viruses

Three multiparous Holstein-Friesian cows late in lactation were inoculated by the intramammary route with 2 mL (0.5 mL per teat) of a virus suspension of H5N1 B3.13 (US-group), containing 10^5.9^ TCID_50_. Three additional animals were similarly inoculated with 10^6.1^ TCID_50_ per 2 mL (0.5 mL per teat) of a virus suspension of H5N1 euDG (EU-group)^21^. One animal was inoculated with 2 mL NaCl and served as negative control (Fig. 1A).

Intramammary inoculation induced clinical disease as early as 1 dpi with impaired general condition, postural abnormalities, and lethargy. All six inoculated cows developed fever (> 40°C) starting at 2 dpi, further exceeding 40.5°C in both groups (Fig. 1B). Moreover, drastically reduced feed intake was observed in both groups of H5N1-infected cows (Fig. 1C). One cow per group (#47 US and #72 EU) displayed clinical signs that met criteria for immediate humane euthanasia at 3 dpi. These included postural and motion disorders, refusal of feed and water intake, dehydration, and severe lethargy. For direct comparison, the control cow (#80) was also euthanized at 3 dpi. Over the next days (#88 EU at 9 dpi and #92 US 13 dpi,), one additional cow from each group deteriorated into clinical conditions meeting humane endpoint criteria (severe lethargy, postural instability, staggering, and signs of respiratory distress).

Prior to infection, daily milk production from individual cows ranged from three to fifteen liters (Fig. 1D). After infection, milk yields rapidly decreased by more than 90%, with only partial recovery observed in the animals remaining at 21 dpi (recovery <3% in #87 US, and up to max. 25% in #66 EU) (Fig. 1D). Starting at 2 dpi, the milk became mucilaginous and viscous and rapidly separated into a serous and a solid fraction with visible curds (Extended Data Fig. 2 A-B). Milk yields in the control animal were low prior to infection, likely due to drying off of this cow (involution in 3 quarters) (Fig. 1D). Onset of severe mastitis was confirmed by California Mastitis Test (CMT) performed daily in both groups (Extended Data Fig. 3, 4, and 5A-C). Clearly positive CMT in infected animals was seen as early as 1 dpi (Extended Data Fig. 4A-C, Extended Data Fig. 5A-C).

In conclusion, calves inoculated oronasally presented signs of mild respiratory illness including nasal mucus secretions and coughing although these cannot be fully associated with outcomes of H5N1 inoculation, whereas intramammary infection of dairy cattle with both clade 2.3.4.4b isolates resulted in severe clinical disease in both cow groups requiring early euthanasia in some cases. Severe disease in lactating cows was accompanied by a drastic reduction in milk production and obvious changes in milk quality.

### Dynamics of viral shedding in calves and cows following HPAIV H5N1 infection

Shedding of IAV RNA was observed in five of six principal-infected calves for a maximum of 8 days, primarily in nasal swabs (Fig. 2A). Generally, low to medium levels of IAV RNA were detected, with peak shedding occurring in nasal swabs between 5-7 dpi in three principal infected calves. IAV RNA was also detected in oral swabs, most frequently at 4-7 dpi, and only seldomly detected as suspect-positive (Cq ≥ 35, single-replicate positive) in rectal swabs. Vaginal and penile swabs, conjunctival swabs collected at necropsy, as well as urine and whole blood, were negative for IAV RNA throughout the study period. All clinical samples collected from sentinel calves were negative for IAV RNA except for two suspect-positive rectal swabs, attributed to environmental contamination during sample collection, suggesting that no transmission of IAV to sentinel calves occurred throughout the study period. Virus isolation and titration were attempted on samples with Cq ≤ 36 (Fig. 2A, Extended Data Table 2). Successful recovery of virus was achieved primarily from nasal swabs of three different calves at 1, 2, 5, and 7 dpi. Titers range from 4.64×10^1^ to 1.7×10^3^ TCID_50_/mL (Fig. 2A).

**Fig. 2.**
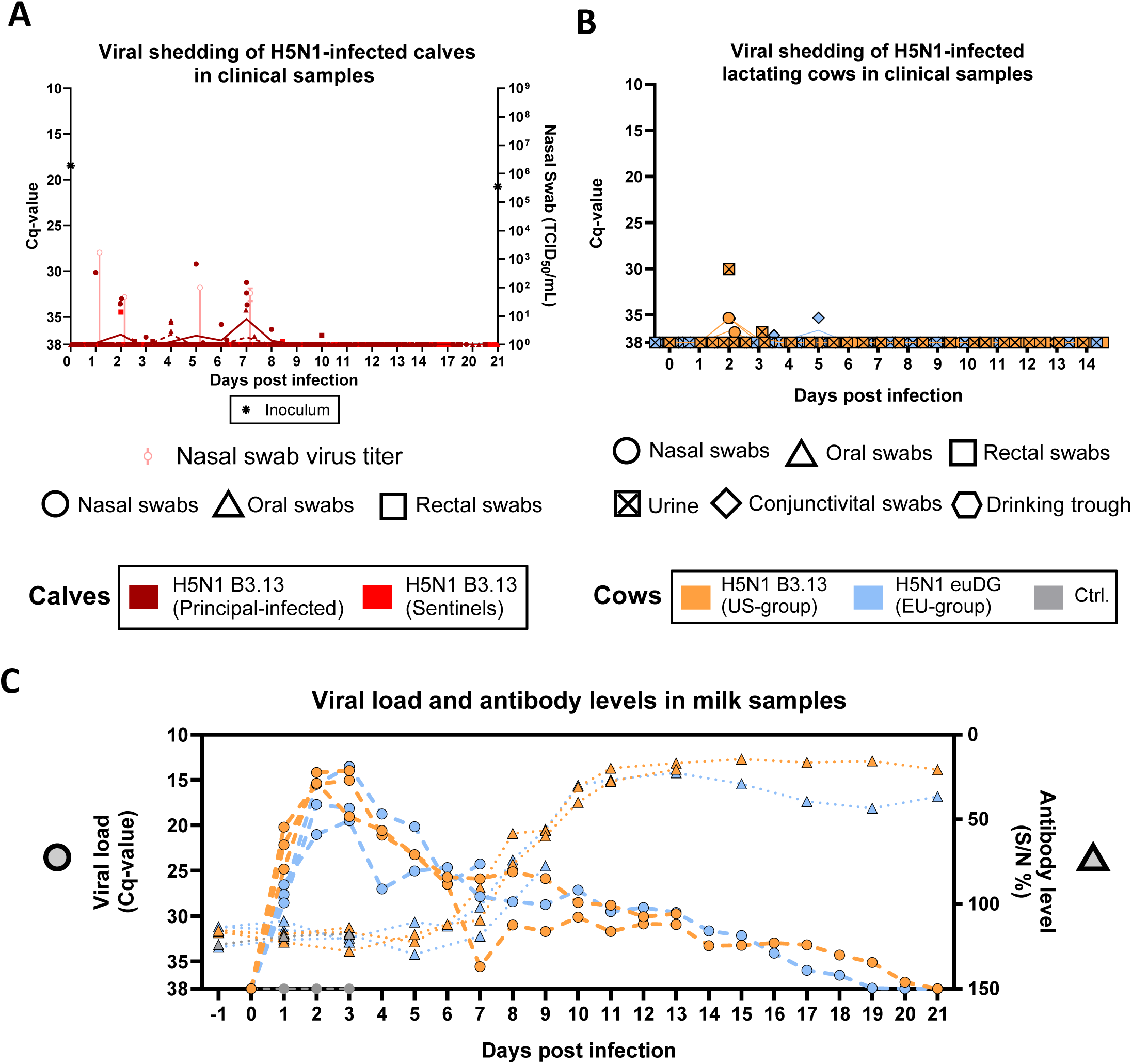
| Viral shedding of influenza A/H5N1 clade 2.3.4.4b virus isolates in experimentally infected calves and lactating cows. **A** RT-qPCR was used for the detection of influenza A M gene (left y-axis) in nasal, oral and rectal swabs collected from H5N1 B3.13 oronasally infected calves post-inoculation an sentinel calves (dark red: principal-infected; bright red: sentinel). Viral titers of nasal swabs (right y-axis) are represented as the mean and standard deviation of nasal swabs with 46.4 TCID_50_/mL on each day. The Cq-value and titer of inoculum are also shown (*) on the respective y-axis. **B** RT-qPCR of nasal, oral, rectal and conjunctivital swabs of lactating cows intramammary infected with H5N1 B3.13 or H5N1 euDG. Orange: Cattle infected with the H5N1 B3.13. Blue: Cattle infected with H5N1 euDG. Grey: Uninfected negative control **C** H5N1 viral genome load (left y-axis) and corresponding H5-specific antibody titers (right y-axis) in milk samples over time. All cattle were milked daily and individual pooled milk samples were analyzed via RT-qPCR for the detection of H5N1 viral RNA. Detection of H5-specific antibodies in selected milk samples was achieved using an H5-specific ELISA and reported as sample OD_450_ / negative control OD_450_ percentage (S/N%).

In lactating cows, RT-qPCR analyses of nasal, conjunctival, rectal swabs and urine samples of animals from the US and EU group, were negative for viral RNA except for two nasal swab samples of #87 and #92 US (Cq value 35 and 36) and two urine samples of #47 and #92 US at 2 and 3 dpi (Cq-values of 30 and 36) (Fig. 2B). In contrast, milk samples showed that all animals in both groups were positive starting at 1 dpi with peak viral genome loads in milk samples at 3 dpi, revealing Cq values ranging from 13 to 21 (Fig. 2C). Viral RNA was detectable in milk samples until 20 dpi and by 9 dpi, antibodies directed against H5 were present in milk samples of each animal and raised to a maximum on 11 dpi and maintained at this level until the end of the study period (Fig. 2C, Table 1).

**Table 1.**
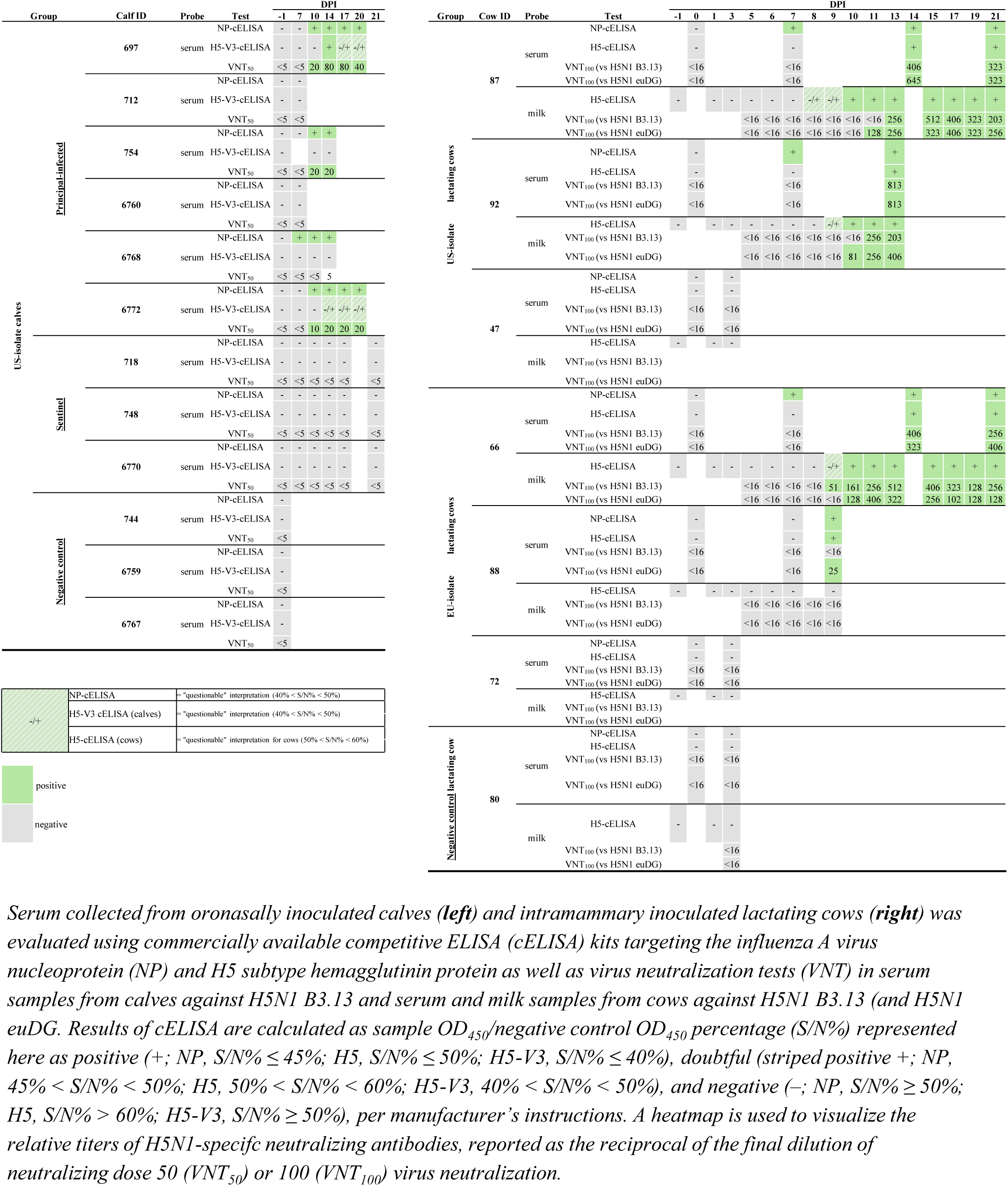
| Influenza A virus-specific serological response in calves and lactating cows following inoculation with HPAIV H5N1 clade 2.3.4.4b.

Virus titration from milk samples was performed from 4 – 13 dpi, but was only successful from 4 – 8 dpi with peak titers of 10^8^ TCID_50_ per mL (Fig. 3A). Due to the milk composition, accurate determination of virus titers from 9 dpi onwards was not possible. Virus titration from mammary gland samples reached peak titers of ∼10^6^ TCID_50_/mL in animals euthanized at 3 dpi (Fig. 3B). Nevertheless, infectious virus could still be isolated from animals euthanized at 9 or 13 dpi (Fig. 3B). Sequencing of viral RNA from mammary gland tissue and milk samples revealed emergence of PB2 amino acid substitution E627K in all three animals after infection with the H5N1 euDG (pooled milk sample of #66 EU day 4 = 20 % K, pooled milk sample of #88 EU day 4 = 20 % K, udder organ sample of #72 day 3 = 89 % K at PB2 position 627) (Fig. 3C). Detection of minor variants at this position from samples of pooled milk from day 1 to day 4 post infection showed that this mutation was acquired early after infection, as it was not detected in the inoculum (Fig. 3C). In the H5N1 B3.13 infected cows, in contrast, marker mutation PB2 M631L was maintained and PB2 E627 remained unaltered (Fig. 3C).

**Fig. 3.**
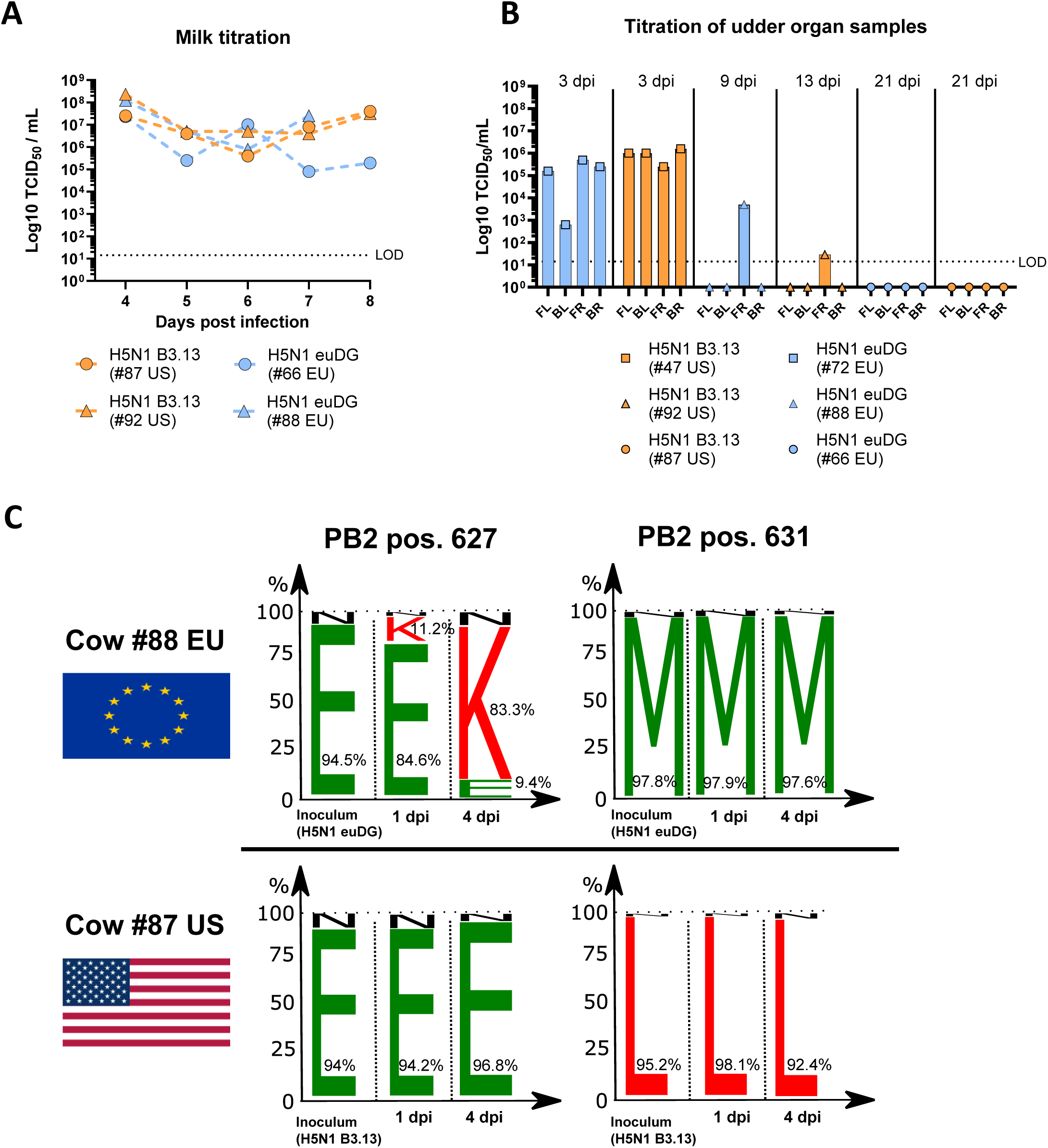
| Infectious virus yields in milk and udder tissue of H5N1 infected lactating cows and genetic adaptations over time. **A** H5N1 viral titers recovered from milk samples of H5N1 B3.13 (orange) and H5N1 euDG (blue) infected lactating cows throughout the experiment. **B** Experimental titration of H5N1 infectious viral particles from individual udder quarters (FL = front left; BL = back left; FR = front right; BR= back right) of each H5N1-infected lactating cow harvested at their respective euthanasia timepoints. **C** Genetic adaptation at position 627 and 631 in the Polymerase Basic Protein 2 (PB2) of H5N1 B3.13 and H5N1 euDG. A sequence logo plot displays the relative proportion of amino acids (E –; K-; N –; M –; L –) present at positions 627 and 631 of polymerase basic protein 2 (PB2) for H5N1 B3.13 and H5N1 euDG present in milk sampled from cows #88 EU and #87 US over time.

The high viral loads in milk samples provided an additional opportunity to validate H5-antigen detection by rapid antigen tests (RATs) (Extended Data Fig. 6). Two out of three H5N1 B3.13-infected animals (#87 & #92 US) were positive in an HA1-HA16-specific RAT at 1 dpi (Extended Data Fig. 6A). By 2 dpi, all H5N1-inoculated cows were positive by RAT (Extended Data Fig. 6B-C) and negative at 10 dpi (Extended Data Fig. 6D), which is consistent with increasing antibody levels in milk from 7 dpi on (Fig. 2C).

In summary, oronasal H5N1 B3.13 inoculation in calves resulted in moderate levels of nasal shedding in five of six principal-infected calves for a maximum of 8 days independent of sex without any evidence for transmission to sentinels. In lactating cows, milk samples obtained from both experimental groups contained high infectious viral loads with viral shedding and genomic detection for up to 8 or 20 dpi, respectively, providing the first evidence for susceptibility of dairy cows to two H5N1 clade 2.3.4.4b viruses belonging to different genotypes from separate continents.

### Mammary gland, not respiratory tract is the primary replication site

Of nearly 40 organ samples collected from each oronasally inoculated calf sacrificed at 7, 14 and 20 dpi, IAV RNA was found present only in mucosa-associated lymphoid tissue (MALT; retropharyngeal lymph node, palatine tonsil, nasopharyngeal tonsil, suppuration from palatine tonsil) of one principal infected calf (#712) at 7 dpi (Extended Data Table 1). All other liquid, swab, and tissue samples (visceral and lymphoid tissues), including lung tissues and bronchoalveolar lavage fluid collected from principal-infected and sentinel calves at 7, 14, 20 and 21 dpi were negative for IAV RNA (Extended Data Table 1). Infectious virus was recovered only from the palatine tonsil suppuration collected post-mortem at 7 dpi.

In lactating cows, RT-qPCR analysis of tissues collected from animals euthanized at 3 dpi revealed peak viral RNA loads in mammary glands, with Cq-values of 20 (#47 US) and 25 (#72 EU) (Extended Data Fig. 1A). At 9 (#88 EU) and 13 dpi (#92 US), viral genome loads in mammary glands were slightly lower than in mammary gland samples collected at 3 dpi and were negative at 21 dpi (#66 EU and #87 US) (Extended Data Fig. 1B). H5N1 viral RNA was also detected at low levels in both groups in neuronal and further tissues at 3 dpi (e.g. #47 US: spinal cord, Cq 28; cerebrum, Cq 36; nervus genitofermoralis Cq 31; #72 EU: nervus genitofermoralis Cq 34) (Extended Data Fig. 1C-D). Organ samples of the respiratory tract remained negative in all animals at respective euthanasia timepoints (Extended Data Fig. 1E). However, no significant histological changes or viral antigen were observed in these tissues. Additionally, there was no IAV RNA detected in whole blood or PBMCs collected from either oronasally inoculated calves or intramammary inoculated cows.

In summary, low levels of IAV RNA were detected only in MALT localized with tissues of the upper respiratory tract in one of two oronasally infected calves at 7 dpi. No IAV RNA was detected in regular anti-mortem collections of whole blood or in samples collected post-mortem from organs, lymphoid tissue, or swabs/fluids from principal-infected and sentinel calves, suggesting that a viremic phase of the infection most likely did not occur. Similarly, intramammary infection of lactating dairy cattle with two distinct genotypes of clade 2.3.4.4b viruses remained restricted to the mammary gland and no evidence of systemic spread was observed.

### Pathology of H5N1 infected calves and cows confirm localized infections

Gross pathology for calves is available in Extended Data Fig. 10. Histological changes in calves oronasally infected with bovine H5N1 at 7 and 14 dpi are depicted in Fig. 4. At 7 dpi, there was a suppurative tracheitis in one animal (#6760) with degenerate neutrophils filling the tracheal lumina. The second calf (#712) at 7 dpi displayed discrete foci of fibrinous interstitial pneumonia with fibrin filling regional alveolar spaces and mild numbers of neutrophils, macrophages and lymphocytes expanding alveolar septa and minimal peribronchiolar inflammatory cells of associated terminal bronchioles. No viral antigen was detected by IHC in these samples (Fig. 4B, D, F). At 14 dpi, the bronchioles of one animal (#754) were lined by hyperplastic epithelium, filled with degenerate neutrophils, and partially occluded by papillary projections composed of a core of fibrous connective tissue with few inflammatory cells and lined by bronchiolar epithelium (bronchiolitis obliterans). Bronchioles were also frequently delimited by prominent lymphoid aggregates (BALT hyperplasia), however, no viral antigen was detected.

**Fig. 4.**
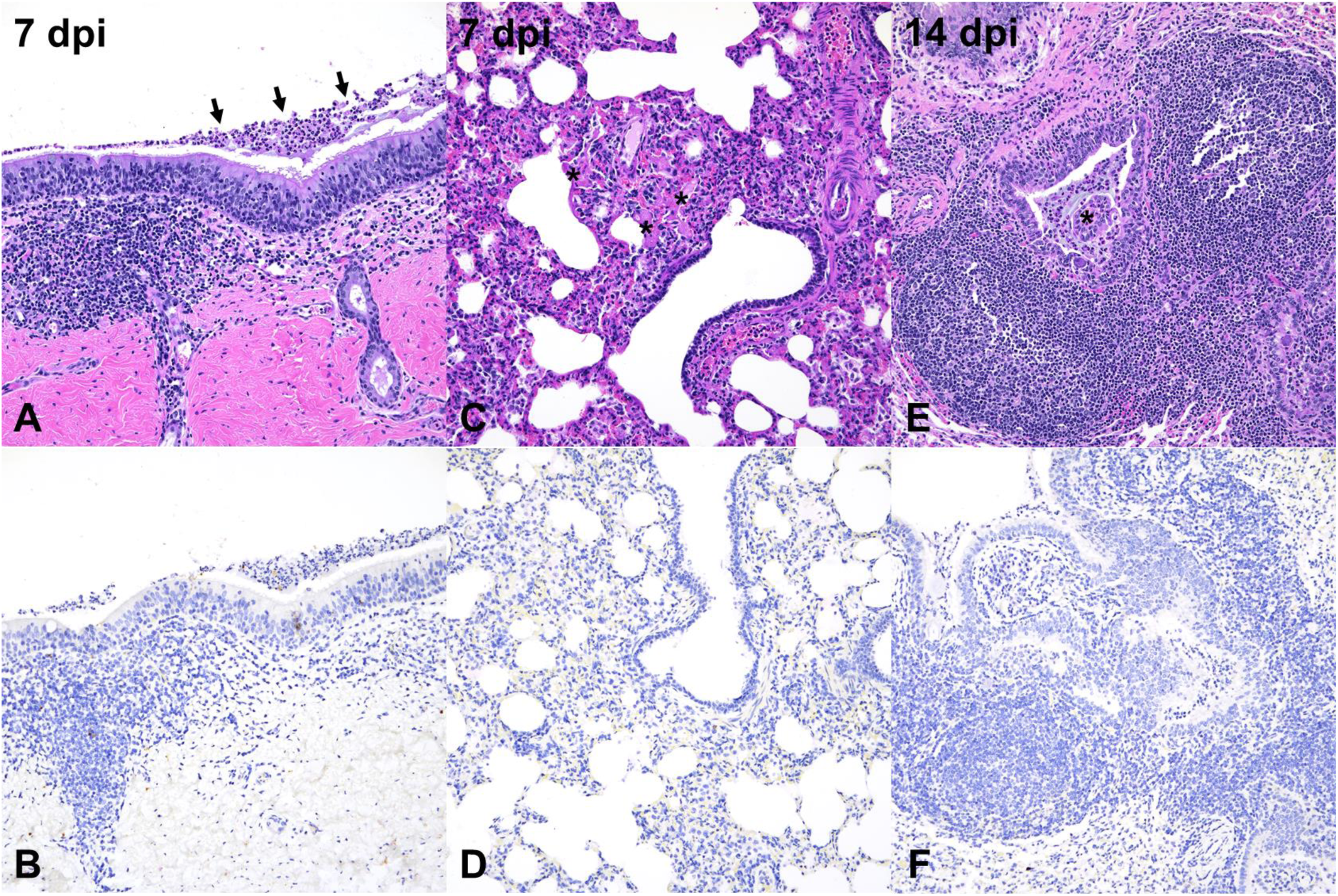
| Histological changes observed in respiratory tissues of calves. Histological changes in calves oronasally infected with H5N1 B3.13. Calves at 7 (A-D) and 14 dpi (E-F) are depicted in the figure. (A and B) H&E staining showing there was a segmental region of suppurative tracheitis at 7 dpi (#6760). Degenerate neutrophils filled the tracheal lumina (arrows). No viral antigen was detected by IHC (B). (C and D) In animal #712 (7 dpi), there were multiple small and discrete foci of interstitial pneumonia (C) with fibrin filling regional alveolar spaces (asterisks) and mild numbers of neutrophils, macrophages and lymphocytes expanding alveolar septa. No viral antigen was detected (D). (E and F) In animal #754 (14 dpi), bronchioles were frequently lined by hyperplastic epithelium, filled with degenerate neutrophils, and partially occluded by papillary projections composed of a core of fibrous connective tissue with few inflammatory cells and lined by bronchiolar epithelium (bronchiolitis obliterans, asterisk). Bronchioles were also frequently delimited by prominent lymphoid aggregates (BALT hyperplasia). No viral antigen was detected (F).

During postmortem examinations of the lactating cows, 45 tissue locations per cow were sampled for histological examination and virus antigen detection (Extended Data Table 5). At 3 dpi, H5N1 B3.13 and euDG induced acute mastitis presented with flocculent material in minimal amounts of milk (#72, EU; #47, US). The character of the histologic changes did not differ between H5N1 B3.13 and euDG. Due to the small number of animals used, a quantitative comparison of the two isolates with regard to the abundance of the virus antigen is not possible (representative pictures included in Extended Data Fig. 7). Up to 90 % of the histological sections of secretory alveoli in the mammary gland evaluated showed acute epithelial necrosis with intraluminal cellular debris admixed with many degenerate neutrophils and intralesional antigen detection (Fig. 5A-B). The basal laminae with lining basal/myoepithelial cells remained largely intact (Fig. 5C). Intralesional virus antigen was confined to the secretory alveolar epithelium and intraluminal cellular debris (Fig. 5D). The teat canal was less prominently affected by necrosis and inflammation, exhibiting IAV nucleoprotein in the remaining lining epithelium (Fig. 5E-F) and debris. The enlarged, draining supramammary lymph node exhibited acute lymphadenitis lacking antigen detection. At 9 dpi (#88 EU), in addition to the acute necrotic lesions, interstitial, mainly lymphocytic infiltrates were present (Fig. 5G) with antigen detection present but limited to up to 50% of the alveoli evaluated histologically, mostly within cellular debris (Fig. 5H) as well as single teat canal lining epithelial cells. At 13 dpi (#92 US), there was still evidence of acute necrosis (Fig. 5I) with scattered antigen detection within cellular debris (Fig. 5J). However, the predominate feature observed at this time point consisted of regenerative and non-suppurative, interstitial inflammatory infiltrates (Fig. 5I). Cellular debris in affected mammary alveoli at 21 dpi still were positive for virus antigen (H5N1 B3.13 only, Fig. 5K). The majority of the examined alveoli were in a regenerative state at 21 dpi (Fig. 5L) with mainly lymphoplasmacytic, interstitial infiltrates (Fig. 5L). All other tissues tested negative for IAV antigen, including those identified to contain low levels of viral RNA in individual animals (spinal cord, cerebrum and genitofemoral nerve, cervix, vestibulum vaginae and the urinary bladder, details included in Extended Fig. 8A-E). The relevance of the intra-or interlobular fibrosis of the inflamed mammary tissue to HPAIV infection could not be established, since it was found in varying degrees in the mammary gland, both in the uninfected control animal (#80, all quarters) and in individual quarters of the infected animals (all infected animals). In accordance with the clinical findings, a lower amount of dry, but otherwise normal ingesta was found in the gastrointestinal tract on 3 and 9 dpi, interpreted a sequela of the HPAIV infection. Additional changes were considered as not being associated with infection but were attributed to the age and lactation status of these multiparous animals (details included in Supplementary Data 1).

**Fig. 5.**
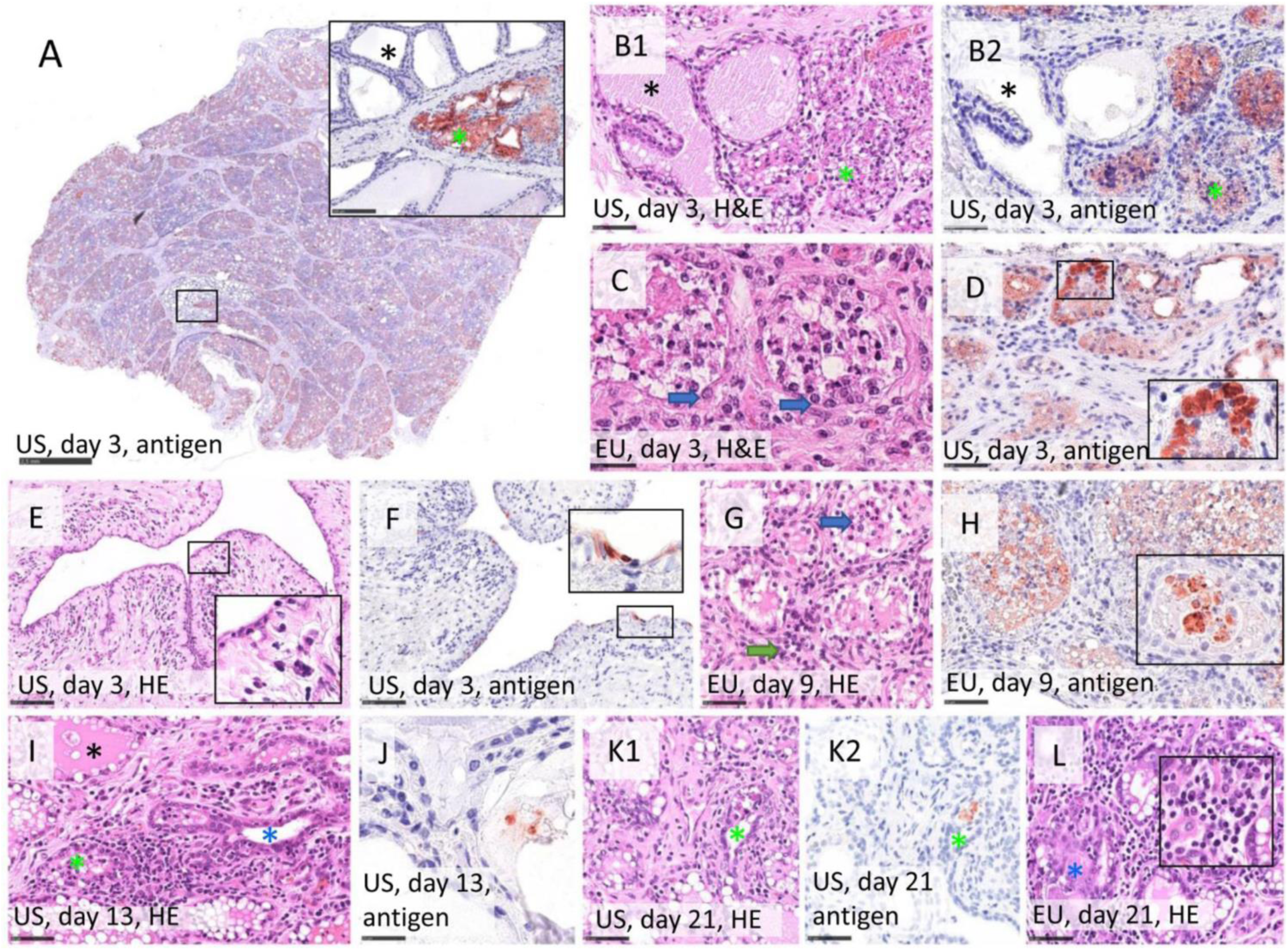
| Histopathology and Influenza A virus (IAV) nucleoprotein (NP) detection in the mammary gland and teat of multiparous cattle after intramammary infection with H5N1 B3.13 and H5N1 euDG. (**A**) Abundant IAV NP detection 3 days post infection (dpi), inlay showing juxtaposition of intact, lactating alveoli lacking antigen (black asterisk) and affected areas (green asterisk) in a lobular pattern. Immunohistochemistry, IHC, using AEC chromogen and Mayer’s hematoxylin counterstain. H5N1 B3.13. Scale bar 2.5 mm and 100 µm (inlay). (**B1**) Full necrosis of the alveolar epithelium with cellular debris filling the lumen and (**B2**) intralesional detection of IAV antigen (green asterisk) on a consecutive slide. Some adjacent alveoli remain unaffected (black asterisk), 3 dpi, H5N1 B3.13. HE (**B1**) and IHC (**B2**). Scale bar 50µm. (**C**) Alveoli affected by necrosis with mostly intact basal lamina lined by basal/myoepithelial cells (blue arrow), indicative for regenerative capacity, 3 dpi, H5N1 euDG. HE. Scale bar 25µm. (**D**) Target cells identified based on morphology following IHC included alveolar secretory epithelium, 3 dpi, H5N1 B3.13. IHC. Sale bar 50µm. (**E**) Teat with diffuse degeneration and necrosis of the lining epithelium, subepithelial edema and mainly neutrophilic infiltrates (inlay), 3 dpi, H5N1 B3.13. HE. Scale bar 100 µm. (**F**) Target cells identified based on morphology following IHC included teat canal epithelium (inlay), 3 dpi, H5N1 B3.13. IHC. Scale bar 100 µm. (**G**) Necrotic alveoli filled with cellular debris admixed with degenerate neutrophils (blue arrow) in acute lesions and many lymphocytes, fewer macrophages, neutrophils and plasma cells in the interstitium (green arrow), H5N1 euDG, 9 dpi. HE. Scale bar 50 µm. (**H**) Abundant intralesional IAV NP detection in secretory alveoli, mainly within cellular debris (inlay), 9 dpi, H5N1 euDG. IHC. Scale bar 50 µm. (**I**) Simultaneous occurrence of either intact, lactating alveoli (black asterisk), disruption of alveolar epithelium by necrosis (green asterisk) and beginning regeneration (blue asterisk), 13 dpi H5N1 B3.13. HE. Scale bar 50 µm. (**J**) Late stage IAV NP detection restricted to cellular debris, found scattered at 13 dpi, H5N1 B3.13. IHC. Scale bar 25 µm. (**K1**) Mammary alveolus with intraluminal sloughed epithelium and cellular debris (green asterisk), and (**K2**) intralesional detection of IAV antigen on a consecutive slide (green asterisk), 21 dpi, H5N1 B3.13. HE (**K1**) and IHC (**K2**). Scale bar 50µm. (**L**) Regenerating alveoli (blue asterisk) with lack of IAV antigen labeling (not shown). Interstitial immune cell infiltrates constitute many lymphocytes and plasma cells (inlay), 21 dpi, H5N1 euDG. HE. Scale bar 50 µm.

### Rapid seroconversion in directly inoculated calves and lactating cows

IAV-specific antibodies present in the serum collected from oronasally inoculated calves was first evaluated using an NP-specific cELISA. All four principal-infected calves were seropositive by 10 dpi (Table 1). Subsequent evaluation of serum using the H5-subtype specific cELISA resulted in just two calves seropositive by 14 dpi. However, neutralizing antibodies were detected in three of four principal-infected calves remaining at 10 dpi with all four becoming positive by 14 dpi (Table 1). Neutralizing titers ranged from 1:5 to 1:80 and were maintained in calves until they were humanely euthanized or on 14 or 20 dpi (Table 1). Only three calves were seropositive (by 14 dpi) when serum was evaluated with the H5-specific c-ELISA. There was no IAV-specific antibody response detected in the sentinel or negative control calves prior to or throughout the study period (Table 1).

Influenza A virus-specific antibodies of infected lactating cows were analyzed by ELISA and virus neutralization tests (VNT) in serum and milk samples throughout the experiment. At 7 dpi, serum samples from 2/2 cows inoculated with H5N1 B3.13 and 1/2 cows inoculated with H5N1 euDG were positive in both the NP-specific and the H5-specific ELISAs (Fig. 2C, Table 1). Neutralizing antibodies in milk samples of infected cows against H5N1 clade 2.3.4.4b were detectable from 9 dpi on and ranged from 1:25 to 1:813 (Table 1). IAV specific antibody detection in the milk was roughly two days delayed in comparison to sera (Table 1).

## Discussion

H5N1 B3.13 is the first influenza A virus reported to circulate efficiently among dairy cattle with widespread dissemination between US farms and onward transmission to various avian and mammalian species, including humans^9,17^. This highlights the promiscuous nature of AIVs, greatly expanding the range of potential hosts and clearly demonstrating their potential to spill-over and adapt to new environments. Nevertheless, the capacity for cows to serve as a host supportive of productive HPAIV infection is surprising, as previous experiments have shown a very low susceptibility of young calves to intranasal inoculation with HPAIV H5N1^12^. With this study, we present detailed data on cattle susceptibility to different HPAIV H5N1 genotypes within the broadly circulating clade 2.3.4.4b, providing insights into pathogenesis, potential transmission routes and mammalian adaptation. We demonstrate that high-dose oronasal infection of clinically healthy calves with an early H5N1 B3.13 isolate of an infected US dairy cow showed low to moderate replication confined to the upper respiratory tract and low to moderate oronasal viral shedding, presenting clinically with only mild signs, yet IAV-specific antibody production from 7 dpi onwards. However, modest replication and shedding of principal-infected calves were not sufficient to infect direct contacts despite recovery of live virus from some of these samples through 7 dpi. Interestingly, in a previous study from 2008 at FLI using a mammalian H5N1 isolate from 2006, one sentinel calf seroconverted upon H5N1-infection^12^. Therefore, our results provide evidence that systemic spread or replication in the respiratory tract and transmission to sentinels is limited in oronasally inoculated calves under our experimental conditions. As such, gs/GD H5 pathogenicity in male and non-lactating female calves seems to remain unchanged for the past decades. However, not evaluated here is e.g. the interaction between suckling calves and cows, and the reciprocal transmission that could occur at this interface.

Recently, Baker and colleagues reported results in four heifers inoculated by an aerosol respiratory route. Similarly, as described here, clinical disease was mild and infection was also confirmed by virus detection, lesions, and seroconversion. However, in these aerosol-infected heifers, transmission was not evaluated, but viral antigen was detected in the lung^22^. In our study, pathological alterations in the respiratory tract were limited in H5N1 B3.13 oronasally-infected calves. While the changes noted at 7 dpi and 14 dpi (one animal each) could be consistent with an acute to late stage of IAV infection, lack of intralesional antigen detection precluded its unequivocal confirmation. However, the window for detection of IAV antigen is usually narrow following infection and possibly occurred at low abundance in these calves based on their limited susceptibility. This likely explains the lack of detection of viral antigen at 7 dpi. The overall histologic alterations and their distribution in both animals are not consistent with bacterial bronchopneumonia associated with the bovine respiratory disease complex (BRDC) despite detection of *Pasteurella multocida, Bibersteinia trehalosi*, and *Mannheimia haemolytica* by PCR with high Ct values.

Drastically different outcomes were observed following direct intramammary inoculation of lactating dairy cows with either H5N1 B3.13 or H5N1 euDG. Acute presentation of severe clinical signs including lethargy, fever and impaired general condition were accompanied with abrupt reduction of feed intake and clinical mastitis with immediate and persistent milk losses of more than 90% in all animals, irrespective of the H5N1 virus isolate used. Histopathology identified a severe, acute, diffuse, necrotizing mastitis with intralesional virus antigen detection in the secretory epithelium and teat canal lining epithelium, but no evidence of systemic spread of the pathogen. In our study, humane euthanasia of four individual cows was required prior to the initially planned end points due to the severity of clinical symptoms (Extended Data Fig. 1F). Although clear evidence of increased mortality events in dairy farms across the USA is lacking right now, a recent field observation study demonstrates that two dairy farms reported mortality associated with H5N1-infections^9^. Conversely, the severe clinical presentation observed in our study may also be related to the age and late lactation phase of these cattle (4-8 years of age, just before dry-off). It is possible that confounding co-morbidities common in older dairy cows contributed to the severity of the disease, and that H5N1-associated disease may be milder in younger, monoparous cows at a different stage of lactation or when individual udder quarters are affected.

It also seems likely that udder manifestation is not a unique feature of genotype B3.13 only, but rather a particular intrinsic ability of the bovine udder to be readily susceptible to H5N1 clade 2.3.4.4b viruses as similar tropism and disease were demonstrated in this study with H5N1 euDG. Susceptibility of cattle to IAVs other than H5 is speculative, but there are historic reports of productive intramammary infection in dairy cows by an ancestral human H1N1 (A/PR8)^23–25^. Mammary infections of cattle by AIVs are supported by the recently described α2.3 receptor expression in the udder tissue, but not in the upper respiratory tract of cattle^18^. High levels of infectious virus were isolated only from milk and udder samples of H5N1-infected lactating cows, clearly demonstrating that replication of H5N1 clade 2.3.4.4b is restricted to the mammary glands after intramammary inoculation. In addition, RT-qPCR analysis of environmental samples collected daily from a communal water trough, urine samples, and nasal, conjunctival and rectal swabs from infected lactating cows revealed only trace amounts of viral RNA, indicating that non-milk related transmission routes seem less relevant; however, modes of transmission in adult cows remain to be evaluated in more detail. Based on field reports, cow-to-cow transmission is most likely driven by the milking process, appears to be equipment-related and thus represents a mechanical and anthropogenic event^9,14,17,26^. Milk and milking procedures, in this respect, seems to be the central mediator of spread within holdings. This is further supported by recent research showing that raw milk spiked with HPAIV H5N1 remained infectious on milking machines for several hours^27^ and can be detected in environmental samples collected from a milking parlor^26^.

Furthermore, substitutions in the PB2 gene sequence, such as the M631L mutation in H5N1 B3.13, appear to be very favorable for replication in the mammary gland^14^. We found a similar PB2 adaptation, namely E627K, for H5N1 euDG in our lactating cows, starting already 1 dpi as a minor variant with significant presence of 627K by 3 dpi. Independent appearance of this substitution in 3 out of 3 cows suggests a strong bottleneck and high evolutionary pressure towards this adaptation. Conclusively, PB2 adaptation to mammalian hosts (either at position 627 or 631) seems to be beneficial for mammary gland replication, as these phenotypes remain stable in any tested milk and tissue sample. Interestingly, the H5N1 B3.13 has also acquired the E627K mutation upon replication in a human case^28^ and was present as a minor variant (2%) in the quasi-species in environmental samples from a dairy farm in Kansas^26^. It remains to be determined whether strain– and/or host dependencies drive the selection of the two PB2 mutations and whether they resemble similar phenotypes.

Spill-back of bovine H5N1 B3.13 into multiple poultry farms has been reported^9^, and this may result in increased environmental contamination in poultry, furthering spill-back into wild bird populations. This has recently been proven by an outbreak of H5N1 B3.13 in a large commercial layer chicken farm in Colorado, USA, where several farm workers tested positive for H5N1 after culling the infected animals^29^. The possibility for non-lactating cattle to serve as a virus source for onward transmission to adult dairy cows, poultry or mammalian species such as felines, should be considered as we observed nasal shedding of infectious virus for 7 days. The same scenario also exists for environments contaminated with milk from H5N1-infected cows.

A tailored surveillance strategy is crucial for effective control. We demonstrate here that aside of RT-qPCR, RATs provide a simple testing tool for milk from individual animals and are suitable for the detection of H5N1 clade 2.3.4.4b, including genotype B3.13. Influenza mastitis should be considered as a differential diagnosis whenever milk characteristics change. However, as antibody levels in milk increase and viral loads decrease, antigens will not be detected by RATs anymore, as has been seen in our study. On the other hand, infectious yields in the excretion also drop at this point due to the neutralizing activity of secreted antibodies. The unique situation, that both viral shedding and neutralizing antibody shedding occur only in milk, may also be a great opportunity to control this epidemic and reduce infectious yields in pool milk from affected herds. In addition to genomic surveillance, also serologic surveillance of pool milk from individual herds may be appropriate to assess the distribution of IAV among dairy herds and facilitate control efforts.

In conclusion, we demonstrated that: (i) the H5N1 B3.13 has only a moderate capacity for respiratory replication in young calves and (ii) was not transmitted to sentinel calves, (iii) dairy cattle are readily susceptible to two distinct and geographically-separated H5N1 clade 2.3.4.4b isolates of mammalian or wild-bird origin following intramammary inoculation and (iv) demonstrate tissue-specific efficient replication and milk shedding (v) the clinical picture of severe disease in dairy cattle was identical for both strains with severe mastitis, (vi) high-titer infectious virus is shed in milk for at least ∼8 days. Finally, the manifestation and main replication site for H5N1 B3.13 following intramammary inoculation is the mammary gland, and systemic spread and infection of other organ systems, including the respiratory tract, have not been observed in lactating cows.

Fortunately, no human-to-human transmission has been reported so far, supporting the concept that these strains have not yet overcome critical barriers to enable human-to-human transmission, such as improved receptor binding, pH stability, and MxA escape^30^. The frequent interface between humans and affected animals (cattle, poultry or wild birds) provide opportunities for reassortment of the bovine B3.13 with human seasonal influenza viruses or other AIVs in circulation. Thus, effective mitigation strategies must be urgently outlined and implemented to i) prevent continuous replication and spread of this pathogen in cattle, ii) avoid any further mammalian adaptation, and iii) stop spillover/spillback infections to other livestock, wild birds, other mammals, and humans. Focused efforts to better elucidate transmission pathways and H5N1 ecology in the dairy industry worldwide are critically needed.

## Methods

### Ethics statement / Biosafety

All experiments conducted at Kansas State University were approved and performed under the Kansas State University (KSU) Institutional Biosafety Committee (IBC, Protocol # 1758) and the Institutional Animal Care and Use Committee (IACUC, Protocol # 4992) in compliance with the Animal Welfare Act. All animal and laboratory work involving infectious highly pathogenic avian influenza virus were performed in biosafety level-3+ and −3Ag laboratories and facilities in the Biosecurity Research Institute (BRI) at KSU in Manhattan, KS, USA. Further evaluation of inactivated samples was conducted in BSL-2 laboratories using enhanced biosafety practices.

The lactating dairy cattle experiment was evaluated by the responsible ethics committee of the State Office of Agriculture, Food, Safety, and Fishery in Mecklenburg–Western Pomerania (LALLF M-V) and gained governmental approval under the registration number 7221.3-2-010/23.

### Virus

The European HPAIV isolate A/wild_goose/Germany-NW/00581/2024 (H5N1), genotype DE-23-11-N1.3 euDG (H5N1 euDG)^21^, was supplied by the German National Reference Laboratory for avian Influenza (Timm Harder) at FLI. It was propagated in embryonated SPF-chicken eggs for 5 days at 37 °C, followed by harvesting the allantoic fluid, which in turn served as the virus stock. In post-inoculation sterility testing of the original undiluted H5N1 euDG virus stock, the marginal presence of Enterococcus spp. was confirmed on sheep blood agar. This was confirmed by NGS-analysis, which revealed a close genetic relationship to *Enterococcus casseliflavus*, a commensal to the normal bacterial flora. There was no evidence that this had any effect on the results obtained in the study.

The North American HPAIV H5N1 isolate A/Cattle/Texas/063224-24-1/2024, genotype B3.13 (H5N1 B3.13, GIS AID accession number: EPI_ISL_19155861) administered to calves and cattle in this study was isolated from the milk of infected dairy cattle in Texas, USA, kindly provided by Dr. Diego Diel, Department of Population Medicine and Diagnostic Sciences, College of Veterinary Medicine, Cornell University, Ithaca, NY, USA^9^. Virus stock was propagated using bovine uterine epithelial cells (CAL-1; In-house) for 3 passages. A single passage on MDCK cells was used to propagate viral stocks for application in virus neutralization assays. The titers of both virus preparations were determined by endpoint dilution titration on MDCK type II cells.

### Cells

FLI Riems: Madin-Darby Canine Kidney (MDCK, RIE-1061) type II cells originating from the Collection of Cell Lines in Veterinary Medicine (CCLV) were used. Cells were incubated at 37°C under a 5% CO_2_ atmosphere. Cultivation medium is composed of a mixture of equal volumes of Eagle Minimum Essential Medium (MEM) (Hank’s balanced salts solution) and Eagle MEM (Earle’s balanced salts solution), 2 mM L-Gln, nonessential amino acids, adjusted to 850 mg L^−1^ NaHCO_3_, 120 mg L^-1^ sodium pyruvate, pH 7.2 with 10% fetal calf serum (FCS) (Bio & Sell GmbH).

Kansas State University: Madin-Darby canine kidney (MDCK) cells were maintained in Dulbecco’s Modified Eagle Medium (DMEM; Corning, Manassas, VA, USA), supplemented with 5% fetal bovine serum (FBS; R&D systems, Flower Branch, GA, USA) and 1% antibiotic-antimycotic solution (Gibco, Grand Island, NY, USA). Media used during virus cultivation (VNT, VI) was similar, but deprived of FBS and supplemented with 0.3% bovine serum albumin (BSA; Sigma-Aldrich, Darmstadt, Germany) and 1% minimum essential medium vitamin solution (Gibco, Grand Island, NY, USA), in addition to 1 µg/mL of TPCK-treated trypsin.

## KSU – Calf study

### Experimental design

Twelve Holstein calves (5-6 months of age) of mixed sex were transported from an Iowa livestock operation to Kansas State University College of Veterinary Medicine (KSU-CVM) in Manhattan, KS.

Upon arrival, blood and swabs were collected and screened for current or recent IAV infection as well as various pathogens associated with bovine respiratory disease complex (BRD), including influenza D virus (IDV). Calves were negative for any active or recent infections with IAV based on RT-qPCR and IAV NP-specific cELISA results. Results of the RT-qPCR based BRD panel revealed some calves were qPCR positive for common bovine respiratory pathogens (Extended Data Table 3). Calves were semi-randomly (sorted according to sex; one (female) calf determined to be hermaphroditic; then randomized intro groups) allocated into three experimental groups: principal-infected (n=6); sentinel (n=3); negative controls (n=3). The control animals were humanely euthanized prior to the day of infection (–2 and –3 dpi). Calves were in good health prior to virus infection based on health examinations conducted by KSU veterinarians.

On the day of virus infection, sentinel calves were physically separated and placed up-current of the room’s directional airflow from principal-infected calves. Principal-infected calves were administered a total dose of 1×10^6^ TCID_50_ (5×10^5^ TCID_50_/mL) in 2mL of H5N1 B3.13 applied as follows: 0.5 mL per nostril using an atomization device (MAD Nasal™ atomization device, Teleflex, Morrisville, NC, USA) and 1 mL orally using a syringe. 48-hours post-infection, sentinel calves were co-mingled with principal-infected calves (Fig. 1A).

Clinical observations and rectal temperatures were recorded daily (Fig. 1A). One calf (#6770; sentinel) developed a high fever prior to the start of the experiment, which resolved following treatment with florfenicol. Baseline samples, including swabs and blood, were collected from all calves upon arrival (–8 dpi) and again after an acclimation period outdoors (–1 dpi). Clinical samples, including urine (when possible), nasal-, oral-, rectal-, vaginal/penile-swabs (collected in 2 mL of DMEM containing 1% antibiotic/antimycotic solution) were collected at –1, 1-14, 17, 20/21 days post infection (dpi) (Fig. 1A). Whole blood samples were collected on –1, 1-7, 10, 12, 14, 17, 20/21 dpi (Fig. 1A). Serum samples were collected on –1, 7, 10, 14, 17, 20/21 dpi (Fig. 1A).. Thorough post-mortem examinations were conducted on days 7 (n=2; principal-infected), 14 (n=2; principal-infected), 20 (n=2; principal-infected), and 21 (n=3; sentinel) post infection (Fig. 1A). Apparent gross lesions were documented prior to extensive sampling of tissue as to determine the scope and extent of impacted tissues (tissue tropism) and any correlation with subsequent IAV detected. Lungs were macroscopically evaluated and scored (a report was generated) according to the percent of lung affected (individual lobes and left/right) with gross lesions including congestion with atelectasis or edema, pneumonia, hemorrhage, and plural fibrosis when present (Extended Data Fig. 10; Extended Data Table 4)^31^.

### Lactating dairy cattle study

Seven female multiparous lactating Holstein-Friesian dairy cattle in an age range between four and eight years, at a state of decreasing milk production, and around 12 months after last calving were obtained from a local dairy farm. The animals were kept in three separate animal rooms (3 x 3 x 1 animals per stable) in the BSL-3 animal facility of the Friedrich-Loeffler-Institut, Greifswald – Isle of Riems, Germany. During the 25-day acclimatization period, the animals were milked once per day using a can milking system (Minimelker, Melktechnik-Discount, Bohmte, Germany) and the amount of milk produced per cattle was documented to have a reliable baseline for each individual (Fig. 1A). Additionally, a California-Mastitis-Test (CMT) was performed on each udder quarter from each cattle daily from –1 dpi onwards until each individual endpoint. Briefly, CMT-reagent was mixed with an equal amount of raw milk received directly from the respective cow teat on a special CMT-plate (Extended Data Fig. 5A), followed by gentle swiveling. The CMT was interpreted according to a comparative picture showing either 1) Unchanged color and consistency (negative, –); 2) low mucus formation (altered, +); 3) strong mucus formation (positive, ++) and 4) intense clumpy and gelatinous mucus formation (strongly positive, +++). Milk production percentages were calculated by generating a mean value of –2 to 0 dpi for each individual animal, which was set as 100%. Prior to infection, all animals tested negative for influenza A viral RNA in nasal swabs and milk samples via RT-qPCR, as well as seronegative in an IAV-specific ELISA targeting the viral Nucleoprotein (NP) (ID-Vet). Prior to inoculation, the udder epidermis was cleaned and the teat and teat orifice were disinfected using an alcohol-based disinfectant. A teat drainage cannula was employed to evacuate all residual milk from the cistern and for inoculation into teat a/o gland cistern. Three animals were inoculated intramammary with 10^5.9^ TCID_50_/cattle by equally administering 0.5 mL per teat (∼ 2 mL total volume, 10^5.31^ TCID_50_/teat) of H5N1 B3.13, and three animals were inoculated intramammary with 10^6.1^ TCID_50_/cattle by equally administering 0.5 mL per teat (∼ 2 mL total volume, 10^5.49^ TCID_50_/teat) of H5N1 euDG. One additional animal, kept in a separate unit, served as negative control and received 0.5 mL sodium chloride solution per teat. The infectious virus titers of both inocula were determined by back-titration on MDCK II cells. Cattle were milked daily, and nasal swab samples as well as samples from the drinking trough of the animals were taken daily until 9 dpi (Fig. 1A). Swab samples from conjunctiva and rectal swabs were taken from 4 dpi until 9 dpi (Fig. 1A). Urine was collected until 14 dpi. EDTA blood samples were taken at 1, 3, 7, 10 and 17 dpi (Fig. 1A). Serum samples were generated before inoculation, 7, 14 dpi, and at the day of euthanasia (Fig. 1A).

### Kansas State University: RNA extraction and RT-qPCR analysis for calf experiment

To document the presence of H5N1 infection, clinical samples (swabs, EDTA blood) and clarified 10% (weight:volume) tissue homogenates were combined with equal volumes of RLT lysis buffer (Qiagen, Germantown, MD, USA) prior to total nucleic acid extraction using an automated magnetic bead-based extraction system (Taco Mini™, GeneReach, Taichung City, Taiwan; BioSprint 96, Qiagen, Germantown, MD, USA) in combination with associated reagents (GeneReach, Taichung City, Taiwan), according to previously established protocols^26^. Subsequently, samples were run in duplicate reactions using a one-step RT-qPCR assay targeting the matrix gene segment of IAV employing a modified M + 64 probe and qScript XLT 1-Step RT-qPCR ToughMix (QuantaBio, Beverly, MA, USA) to determine the quantity of IAV RNA, with thermocycling conditions as described previously^26^ A positive Cq cut-off of 38 cycles was established for samples where both wells were positive.

### FLI Riems: Sample collection, RNA extraction and RT-qPCR analysis for lactating dairy cattle study

Raw milk samples were collected individually per quarter and directly used for RNA extraction. In addition, bulk milk samples generated via milking machine were also collected and analyzed. Swabs were taken using rayon swabs (DRYSWAB^TM^ Standard Tips, MWE) and were immediately transferred into 2 mL of cell culture medium containing 1% Baytril (Bayer), 0.5% Lincomycin (WDT) and 0.2% Amphotericin/Gentamycin (Fisher Scientific Waltham). Blood samples were taken using the Kabevette® G system and disposable needles. Organ samples with 2×2×2 mm in size were transferred into a 2 mL collection tube containing 1 mL of cell culture medium containing 1% penicillin-streptomycin (Biochrome) and one stainless steel bead per sample. Homogenization was established by rough shaking of the samples in a TissueLyser II instrument (Qiagen) for 2 min. at 300 Hz. Viral RNA of all samples was extracted using 100 µl of raw milk or sample medium or supernatant in the NucleoMag Vet-kit (Macherey-Nagel) on a BioSprint 96 platform (Qiagen). Detection of H5N1 B3.13 and H5N1 euDG via RT-qPCR was established as recommended and described in the SOP VIR 018 – ed. 02 – 09/23 by the “European Union Reference Laboratory for Avian Influenza and Newcastle Disease” in Italy^32,33^.

### Kansas State University: Virus isolation and virus titration from calve study

Clinical samples and tissue homogenates with Ct values <36 were subjected to virus isolation and virus titration. Viral titration and immunofluorescence assays (IFA) were performed as described previously^31^. Briefly, ten-fold serial dilutions of syringe filtered (0.45µm) samples, tested in four replicates, were transferred onto 96-well plates containing confluent monolayers of MDCK cells. After infecting cells, 96-well plates were incubated at 37 °C and observed daily (light microscope) to monitor the conditions of the cellular monolayer and cytopathic effects/cell morphology. After 48-hours, plates were washed with PBS prior to fixation with ice-cold 100% methanol for 10-minutes at –20°C; then cells were washed with 1x PBS and incubated for 1 hour with influenza A-specific HB65 primary antibody (HB-65; ATCC, Manassas, VA^34^ at room temperature. Subsequent washes with PBS containing 0.05% Tween-20 (PBS-T) were conducted prior to incubation with goat anti-mouse IgG (H+L) secondary antibody (Alexa-488, Fisher Scientific, Waltham, MA) and incubation at room temperature for 30 minutes. Plates were washed and dried prior to observation of fluorescent signal using an EVOS microscope. Viral titers were calculated using the Reed-Muench method^35^ with a limit of detect at 46 TCID_50_/mL.

In parallel, attempts for virus isolation were conducted utilizing 25 cm^2^ flasks (T-25) of confluent MDCK cells. Cells were first washed with 1x PBS to remove any residual FBS and subsequently incubated with 500 µL of diluted (1:10) sample and 2 mL of infection media for 2 hours, gently rocked every 15 minutes, prior to addition of 2.5 mL of infection media. After two-three days of incubation at 37 °C, supernatants were collected from T-25 flasks. Flasks were then subjected to IFA protocols as described above and reported as positive or negative for viral presence based on fluorescent signal.

### Virus isolation and virus titration from cattle study

Virus titers of selected milk and udder organ samples were determined by a TCID_50_ endpoint dilution assay on MDCKII-cells. Briefly, 10-fold serial dilutions of respective samples were prepared and transferred onto 96-well plates containing confluent monolayers of MDCKII-cells (duplicates). Plates were incubated for 72 hours at 37 °C. Virus titer was evaluated by the presence of a specific cytopathic effect (CPE) and was calculated according to the “Measure of Infectious Dose in Specific Infections (midSIN)”-method^36^.

### FLI Riems: Serology of the cattle study

Serological analysis of blood samples or milk samples from all animals was performed by using a commercial IAV-specific enzyme-linked immunosorbent assay (ELISA) detecting NP– and H5-specific antibodies (ID-Vet, Montpellier, France) according to the manufacturer’s instructions.

Virus neutralizing antibodies were investigated via a virus neutralization test (VNT_100_) on MDCK II cells. In brief, serum samples of respective timepoints were serially diluted in DMEM on a 96-well plate (log2 steps) and mixed with 100 TCID_50_ of H5N1 B3.13 or H5N1 euDG followed by incubation for 1h at 37 °C. Subsequently, 100 µl of MDCK II cells were added to each well, followed by another incubation period for 72h at 37 °C. Neutralizing antibodies were evaluated and recognized by light microscopy in the absence of a CPE. The last serum dilution with intact cells and no visible CPE was considered as neutralization titer against the respective virus.

### Kansas State University: Serology and host immune responses

Serum, collected from all calves upon arrival (–8 dpi), prior to infection (–1 dpi), and at defined time points (7, 10, 14, 17, 20/21 dpi) throughout the study, were evaluated for the presence of influenza A specific antibodies using several methods.

Neutralizing antibody titers were determined according to previously established protocols^37^ with slight modifications. Briefly, heat-inactivated serum was combined with an equal volume of H5N1 B3.13 stock virus (propagated additionally once on MDCK cells), diluted to 100 TCID_50_/50 µL, in duplicate wells on 96-well plates and incubated at 37°C for 1 hour. Subsequently, the serum/virus mixture was transferred to 96-well plates containing confluent monolayers of MDCK cells. After 48 hours of incubation, IFA was preformed (similar to described above for virus titration/isolation) and neutralizing antibody titers were recorded as 50% inhibition of virus growth per well.

Additionally, commercially available enzyme-linked immunosorbent assay (ELISA) kits, validated for application with bovine-origin serum, targeting: (i) the conserved IAV nucleoprotein (NP-ELISA; ID Screen^®^ Influenza A Antibody Competition Multi-species, Innovative Diagnostics, France) and (ii) the H5-specific hemagglutinin protein (H5-ELISA; ID Screen^®^ Influenza H5 multi-species competitive ELISA V3, Innovative Diagnostics, France) were used according to manufacturer’s instructions.

### Kansas State University: Macroscopic and microscopic pathology

Post-mortem examinations were conducted at 7, 14 and 20/21 dpi. Macroscopic pathology was determined and scored in toto and the percentage of lung lesions were reported, based on previously published protocols^31^. Tissue samples were fixed in 10% neutral buffered formalin for a minimum of 7 days and subsequently transferred to 70% ethanol and processed using standard histological techniques, and stained with hematoxylin and eosin (H&E). Collections included paired collections (fresh tissue and formalin fixed) of representative sections (and lesions) obtained from respiratory, gastrointestinal, and reproductive tract, lymphoid tissue including spleen, various lymph nodes and musical associated lymphoid tissue, brain, eyes and, eye lids, and other tissues including adrenal gland, heart, and pancreas. Bronchoalveolar lavage fluid (BALF) was collected from the right side of the lung. Fluids collected included aqueous humor, cerebral spinal fluid, and bile. Transudates from the pericardial sac, thoracic cavity and abdominal cavity and urine were also collected when present Conjunctival swabs and swabs from the reproductive tract also collected at necropsy. Selection of animals for necropsy was based on sex to ensure representative samples from each gender, as well as on data that was available regarding viral RNA shedding in nasal and oral swabs. Four-micron sections from the lower respiratory tract tissues were stained with routine hematoxylin and eosin after standard automated processing and paraffin embedding. Tissues were subsequently examined microscopically by a board-certified veterinary pathologist.

### Kansas State University: Influenza A virus-specific immunohistochemistry (IHC)

Immunohistochemistry (IHC) for detection of IAV H5N1 nucleoprotein (NP) antigen was performed on the automated BOND RXm platform and the Polymer Refine Red Detection kit (Leica Biosystems, Buffalo Grove, IL). Following automated deparaffinization, four-micron formalin-fixed, paraffin-embedded tissue sections on positively charged Superfrost® Plus slides (VWR, Radnor, PA) were subjected to automated heat-induced epitope retrieval (HIER) using a ready-to-use EDTA-based retrieval solution (pH 9.0, Leica Biosystems) at 100°C for 20 min. Subsequently, tissue sections were incubated with the primary antibody (rabbit polyclonal anti-Influenza A virus NP (Cell Signaling Technology, #99797/F8L6X) diluted 1:1,200 in Primary Antibody diluent [Leica Biosystems]) for 30 min at ambient temperature followed by a polymer-labeled goat anti-rabbit IgG coupled with alkaline phosphatase (30 min). Fast Red was used as the chromogen (15 min), and counterstaining was performed with hematoxylin for 5 min. Slides were dried in a 60°C oven for 30 min and mounted with a permanent mounting medium (Micromount®, Leica Biosystems). Lung sections from a pig experimentally infected with swine influenza virus A/swine/Texas/4199-2/1998 H3N2 and mink-derived clade 2.3.4.4b H5N1 isolate, A/Mink/Spain/3691-8_22VIR10586-10/2022 were used as positive assay controls (Extended Data Fig. 9).

### FLI Riems: Pathology

Full autopsy was performed on all animals under BSL3 conditions and macroscopic diagnoses were recorded. In total, 45 samples (Extended Data Table 5) were fixed in 10% neutral buffered formalin. Tissues were paraffin-embedded and 2-4-μm-thick sections were stained with hematoxylin and eosin (HE). Consecutive slides were processed for immunohistochemistry according to standardized procedures of avidin-biotin-peroxidase complex-method as described^38^. The primary antibody against the IAV nucleoprotein was applied overnight at 4°C (ATCC clone HB-64, 1:200), the secondary biotinylated goat anti-mouse antibody was applied for 30 minutes at room temperature (Vector Laboratories, Burlingame, CA, USA, 1:200). Color was developed by incubating the slides with avidin-biotin-peroxidase complex solution (Vectastain Elite ABC Kit; Vector Laboratories), followed by exposure to 3-amino-9-ethylcarbazole substrate (AEC, Dako, Carpinteria, CA, USA). The sections were counterstained with Mayer’s hematoxylin and coverslipped. As negative control, the non-infected cow was tested with the primary antibody, and consecutive sections of infected animals were labelled with an irrelevant antibody (anti Sars clone 4F3C4, 1:45)^39^. A positive control slide from an IAV infected chicken was included in each run (details included in Extended Data Fig. 8F-H). All sides were scanned using a Hamamatsu S60 scanner, evaluation was done using the NDPview.2 plus software (Version 2.8.24, Hamamatsu Photonics, K.K. Japan) by a trained pathologist (TB) and a board-certified pathologist (AB, DiplECVP). HE stained sections were evaluated and described. Following IHC the distribution of virus antigen was graded on an ordinal scale with scores 0 = no antigen, 1 = focal, affected cells/tissue <5% or up to 3 foci per tissue; 2 = multifocal, 6%–40% affected; 3 = coalescing, 41%–80% affected; 4 = diffuse, >80% affected. The target cell was identified based on the morphology.

### Kansas State University: Next-generation sequencing (NGS)

Samples with high quality RNA extracts were subjected to previously established next-generation sequencing (NGS) methods^31^ in order to evaluate the presence/frequency of genomic variants and their potential relation to host-adaptation (in reference to/changes compared to inoculum). The whole genome sequence of the cattle –derived clade 2.3.4.4b H5N1 virus was determined using the Illumina NextSeq sequencing platform (Illumina, San Diego, CA, USA). Briefly, viral RNA was extracted from the infection inoculum and VI-positive clinical samples (inactivated in RLT lysis buffer; Qiagen, Germantown, MD, USA) using the QIAamp viral RNA mini kit (Qiagen, Germantown, MD, USA). Viral gene segments for infection inoculum and clinical samples were amplified using SuperScript™ III One-Step RT-PCR System with Platinum™ Taq DNA Polymerase (Thermo Fisher Scientific, Waltham, WA, USA) with a universal influenza primer set^40,41^. All samples were normalized to 20 ng/µl (100-300ng) prior to library preparations. Sequencing libraries were prepared using the Illumina DNA Prep kit (Illumina, San Diego, CA). Libraries were sequenced using pair-end chemistry on the Illumina NextSeq platform with the NextSeq 500/550 Mid Output Kit v2.5 (300 cycles). Sequencing reads were demultiplexed and parsed into individual FASTQ files and imported into CLC Genomics Workbench version 23.0.5 (Qiagen, Germantown, MD, USA) for analysis. Reads were trimmed to remove primer sequences and filtered to remove low quality and short reads. The trimmed reads were mapped to the reference sequence (A/Cattle/Texas/063224-24-1/2024; GISAID: EPI_ISL_19155861). Following read mapping, all samples were run through the low frequency variant caller module within CLC Genomic Workbench with a frequency cutoff greater than 2%.

### MinION sequencing

MinION-based sequencing of avian influenza positive samples with Cq values < 28 was carried out as described before^42,43^. Briefly, the RNA was transcribed into DNA using Superscript 60 III One-Step and Platinum Taq (#12574026, Thermo Fisher Scientific, USA) Kit with influenza-specific primers (Pan-IVA-1F_BsmF (26mer wobbel) TATTCGTCTCAGGGAGCRAAAGCAGG; Pan-IVA-1R_BsmR (26mer wobbel) ATATCGTCTCGTATTAGTAGAAACAAGG). DNA amplificants were purified with Agencourt AMPure XP beads (#A63881, Beckmann Coulter, Krefeld, Germany) magnetic beads using DNA LoBind® Tubes (#0030108051, Eppendorf, Wesseling-Berzdorf, Germany). Approximately 200 ng of DNA per sample was used for sequencing by a transposase-based library preparation approach with Rapid Barcoding (SQK-RBK114.24, Oxford Nanopore Technologies, Oxford, UK) and a PromethION Flow Cell (FLO-PRO114M) on a PromethION 2 solo device with MinKNOW Software Core (v5.9.12). Live high accuracy base calling of the raw data with Dorado (v7.3.11, Oxford Nanopore Technologies) was followed by demultiplexing, a quality check and a trimming step to remove low quality, primer and short (<20 bp) sequences. For analyzing, the bioinformatic software suite Geneious Prime^®^ (Biomatters, Version 2024.0.5) was used. The sequences were trimmed, to remove the primer sequences. Consensus sequences were obtained with an iterative map-to-reference approach with Minimap2 (vs 2.24). The H5N1 B3.13 or the H5N1 euDG isolate sequence was used as a reference. Polishing of the final genome sequences and annotation was done manually after consensus generation (threshold matching 60% of bases of total adjusted quality). A total amount of n=53 samples from the animal experiment as well as both virus stocks were sequenced. For the majority (42 samples) only partial assemblies were achieved. The remaining samples were screened for amino acid exchanges. For major variants A threshold of 55% was used to search for specific mutations in the consensus sequences within the sequences. For adaptive mutations (PB2 E627K) also minor variants were determined.

### IAV-specific Rapid-Antigen-Test (RAT) evaluation

Milk samples from the animal experiment served as samples for validation of this assay with application to bovine milk origin samples. Serial samples of infected animals were analyzed in the HA1-HA16 specific Megacor test. Briefly, the provided swab was dipped into the respective milk sample and afterwards transferred into the assay buffer and mixed according to the manufacturer’s protocol. Afterwards, the test strip provided in the kit was dipped into the assay buffer according to manufacturer’s instructions. Results were read after 15 minutes of incubation.

## Acknowledgements

This work was supported by USDA NACA #58-3022-3-004, National Bio and Agro-Defense Facility (NBAF) Transition Fund from the State of Kansas, the USDA Animal Plant Health Inspection Service’s NBAF Scientist Training Program, the AMP and MCB Cores of the Center on Emerging and Zoonotic Infectious Diseases (CEZID) of the National Institutes of General Medical Sciences under award number P20GM130448, and the NIAID supported Centers of Excellence for Influenza Research and Response (CEIRR, contract number 75N93021C00016). This work was further funded by the DURABLE project, co-funded by the European Union, under the EU4Health Programme (EU4H), Project no. 101102733, and the Kappa-Flu project, under the Horizon Europe Programme (grant agreement KAPPA-FLU no. 101084171) and the German Federal Ministry of Education and Research within the project ‘PREPMEDVET’ grant no. 13N15449.

Invaluable contributions that supported the success of this work were made by the professional and technical associates/staff of KSU-CEEZAD/CEZID personnel (not limited to) Govindsamy Vediyappan, Eu Lim Lyoo, Shanmugasundaram Elango, Shristi Ghimire, Patricia Assato, Daniel Madden, Yonghai Li, Isaac Fitz, and Zane Kohl. Additional support was provided by KSU-VDL molecular diagnostic and histopathology section lab personnel (oversight by Jamie Retallick, Greg Hazlicek, and Lance Knoll), personnel of the histology and immunohistochemistry sections of Louisiana Animal Disease Diagnostic Laboratory at Louisiana State University and the coordination and oversight provided by the animal care, lab coordinators, and biosafety staff of the Biosecurity Research Institute (BRI) at Kansas State University.

We also thank Mareen Grawe, Silvia Schuparis, and Robin Brand for excellent technical assistance. We also thank Frank Klipp, Steffen Kiepert, Christian Lipinski, Felix Zimak, René Siewert, Ralf Henkel, Ralf Redmer, Marco Beerbohm, and Andreas Bath for their invaluable and dedicated animal care. We are grateful to editorial remarks on the text by Timm Harder and scientific advice by Lars Mundhenk and Christian Grund.

We also thank Innovative Diagnostics for kindly providing the H5-ELISA kits.

## Author contributions

Conceptualization: MB, JAR, NJH, LU, DH, AB; Data Curation: NJH, KC, AKA, AP, TB, AB, CDM, JDT, MC, ELL, LU, JS; Methodology: NJH, KC, LU, TB, JDT, MC, TK, FMF, JS, DH, RP, BC, GS, SK, NNG, UBB, LH, IM, MN, LMC; Formal analysis: NJH, KC, AKA, LU, TB, AB, CDM, JDT, MC, AP; Investigation: LU, NJH, TB, AB, KC, JDT, MC, FMF, JS, RP, VPR; Visualization: NJH, JS, AB, KC; Writing – Original Draft: NJH, KC, LU, MB, JAR; Writing – Review and Editing: MB, JAR, DH, LU, NJH, KC, AB, JS, JDT, FMF, AKA, AP, DD; Supervision: MB, JAR, AB, DH, LU; Funding acquisition: MB, JAR

## Competing interests

The J.A.R. laboratory received support from Tonix Pharmaceuticals, Genus plc, Xing Technologies, and Zoetis, outside of the reported work. J.A.R. is inventor on patents and patent applications on the use of antivirals and vaccines for the treatment and prevention of virus infections, owned by Kansas State University. The other authors declare no competing interests.

## Data availability

Consensus sequences of both isolates used for inoculation are available in the INSDC under accession PQ106994-PQ107009 (H5N1 B3.13: PQ106994-PQ107001; H5N1 euDG: PQ107002-PQ107009). Raw data were filed to the SRA under project number PRJNA1141392.

## Extended Data Figures

**Extended Data Fig. 1.**
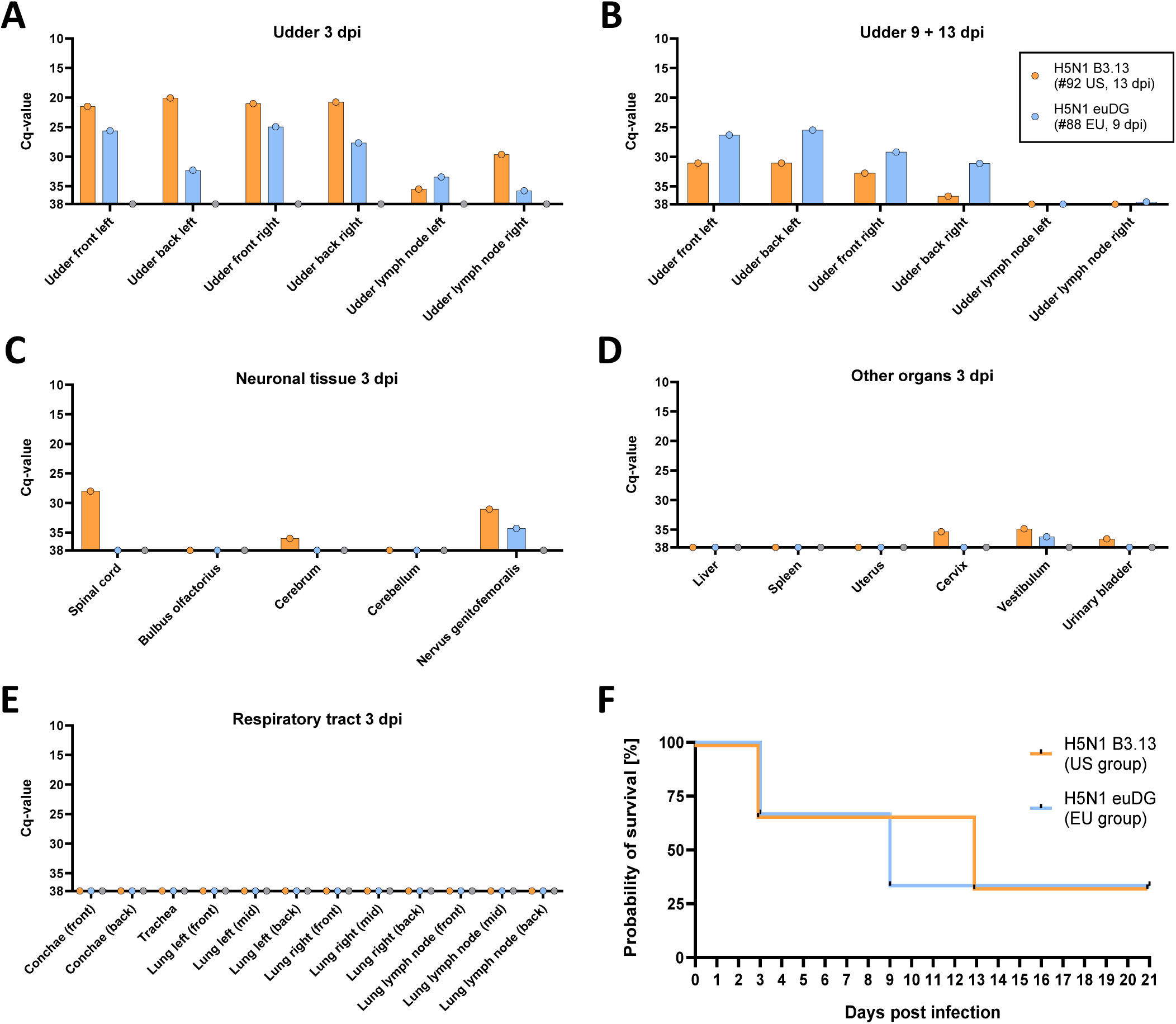
| Viral genome load in organ samples and survival data of lactating cows. Orange: lactating cows infected with H5N1 B3.13. Blue: Lactating cows infected with H5N1 euDG. Grey: Uninfected negative control cow. **A** Viral genome load in udder organ samples of lactating cows euthanized at 3 dpi (#72 EU, #47 US, #80 Ctrl.) **B** Viral RNA load in udder organ samples of lactating cows euthanized at 9 dpi (#88 EU) or 13 dpi (#92 US). **C** Viral genome load in organ samples from neuronal tissues. **D** Viral RNA load in other internal organs of lactating cows. **E** Viral genome load in organ samples of the respiratory tract. **F** Survival curve of lactating cows over the course of the experiment.

**Extended Data Fig. 2.**
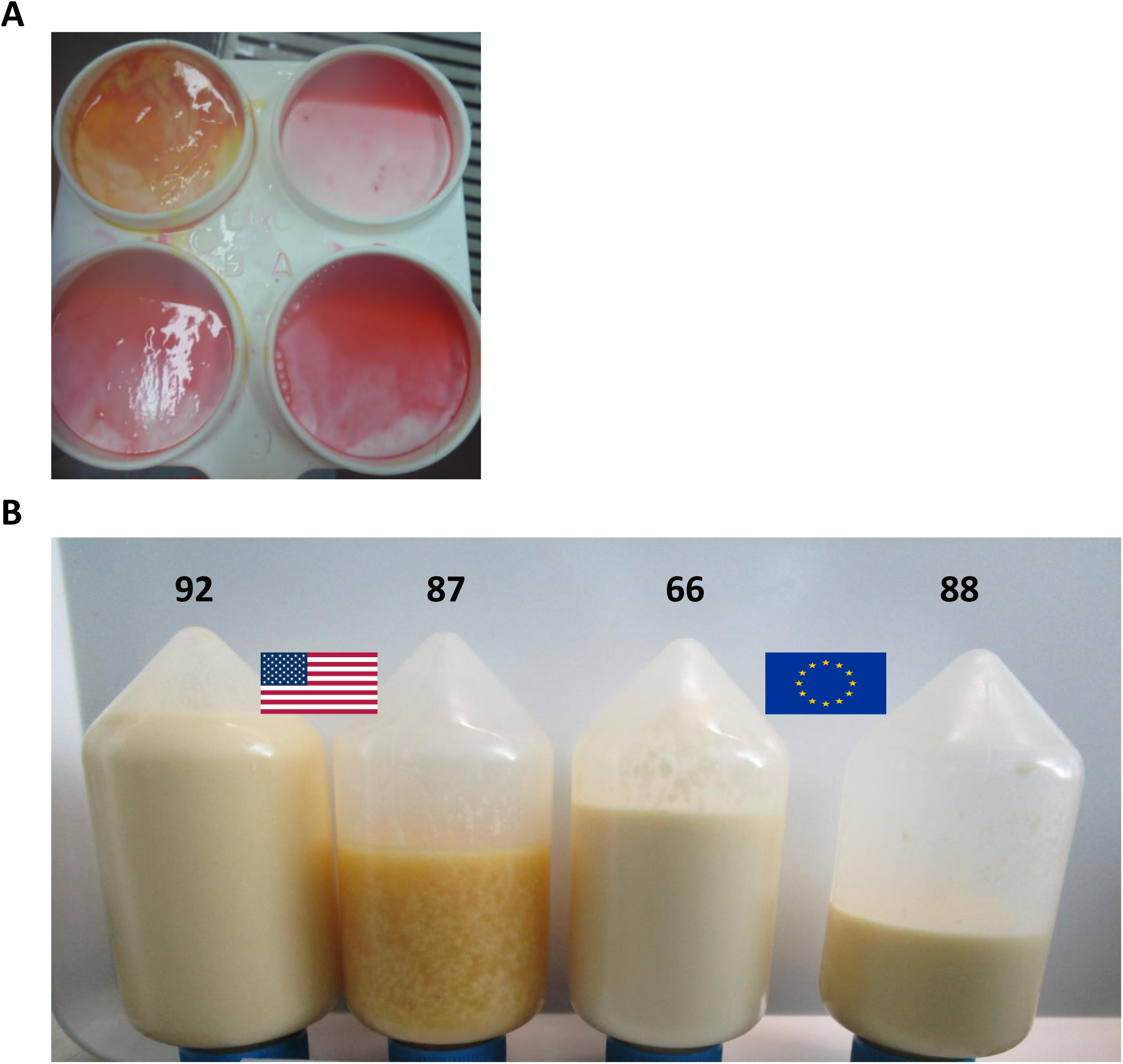
| Exemplary milk consistency and appropriate CMT of H5N1-infected lactating dairy cows during the animal trial. **A** CMT-picture of an H5N1 euDG-infected lactating dairy cow at 2 dpi **B** Milk consistency of H5N1 B3.13 and euDG infected dairy cows during the experiment (4 dpi)

**Extended Data Fig. 3.**
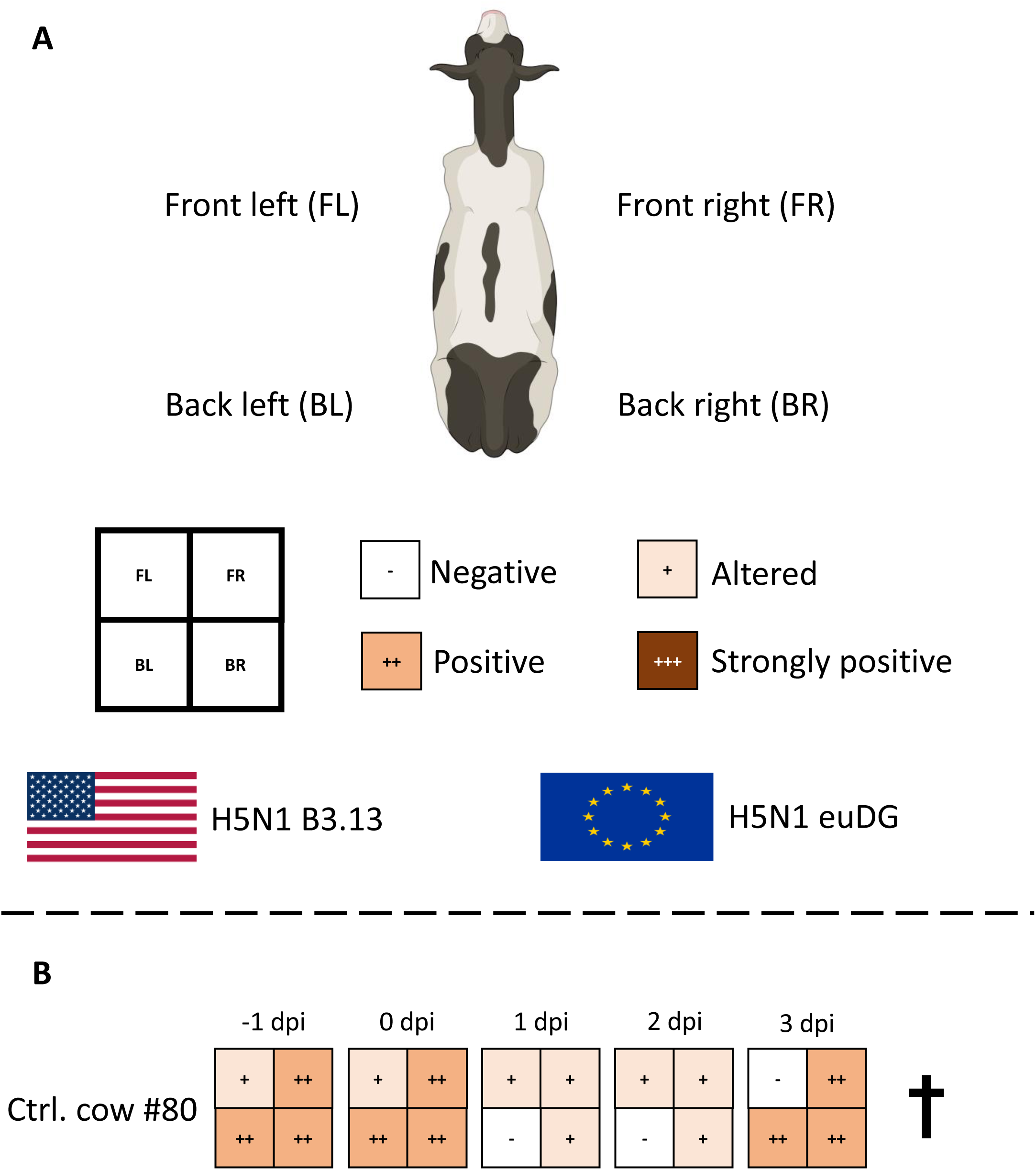
| California Mastitis Test (CMT) **A** Legend for semi-quantification. Milk samples from individual quarters (front left/right and back left/right were gained and collected on appropriate CMT-plates. CMT-reagent was applied ∼ 1:1 to the milk samples and was graded by eye with the help of a defined template. **B** CMT of milk samples from the uninfected control animal (#80) during the course of the experiment until its euthanasia timepoint.

**Extended Data Fig. 4.**
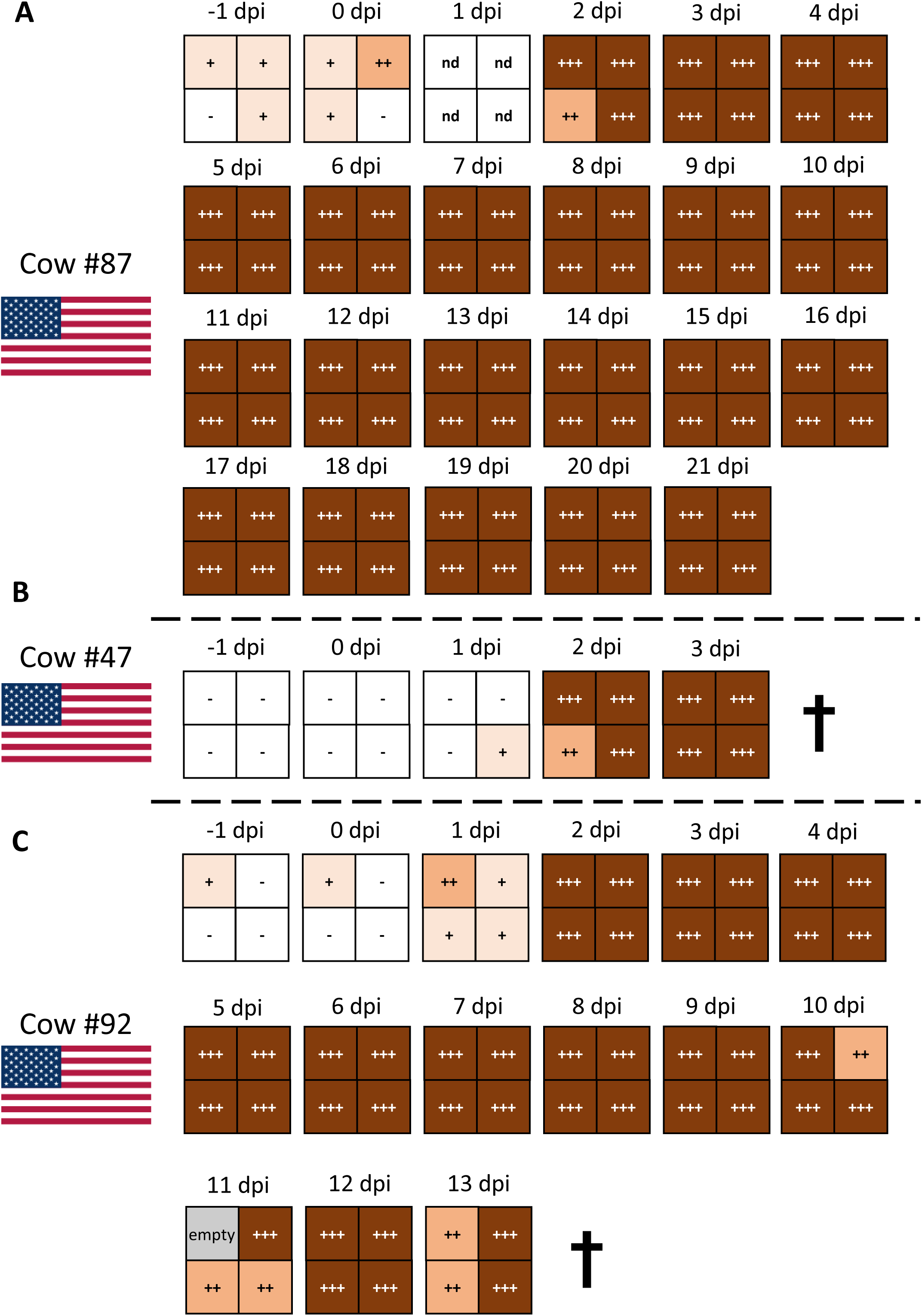
| California Mastitis Test (CMT) of lactating cows infected with H5N1 B3.13 (US-group) **A – C** CMT of milk samples from cattle infected with H5N1 B3.13 during the course of the experiment until their respective euthanasia timepoint.

**Extended Data Fig. 5.**
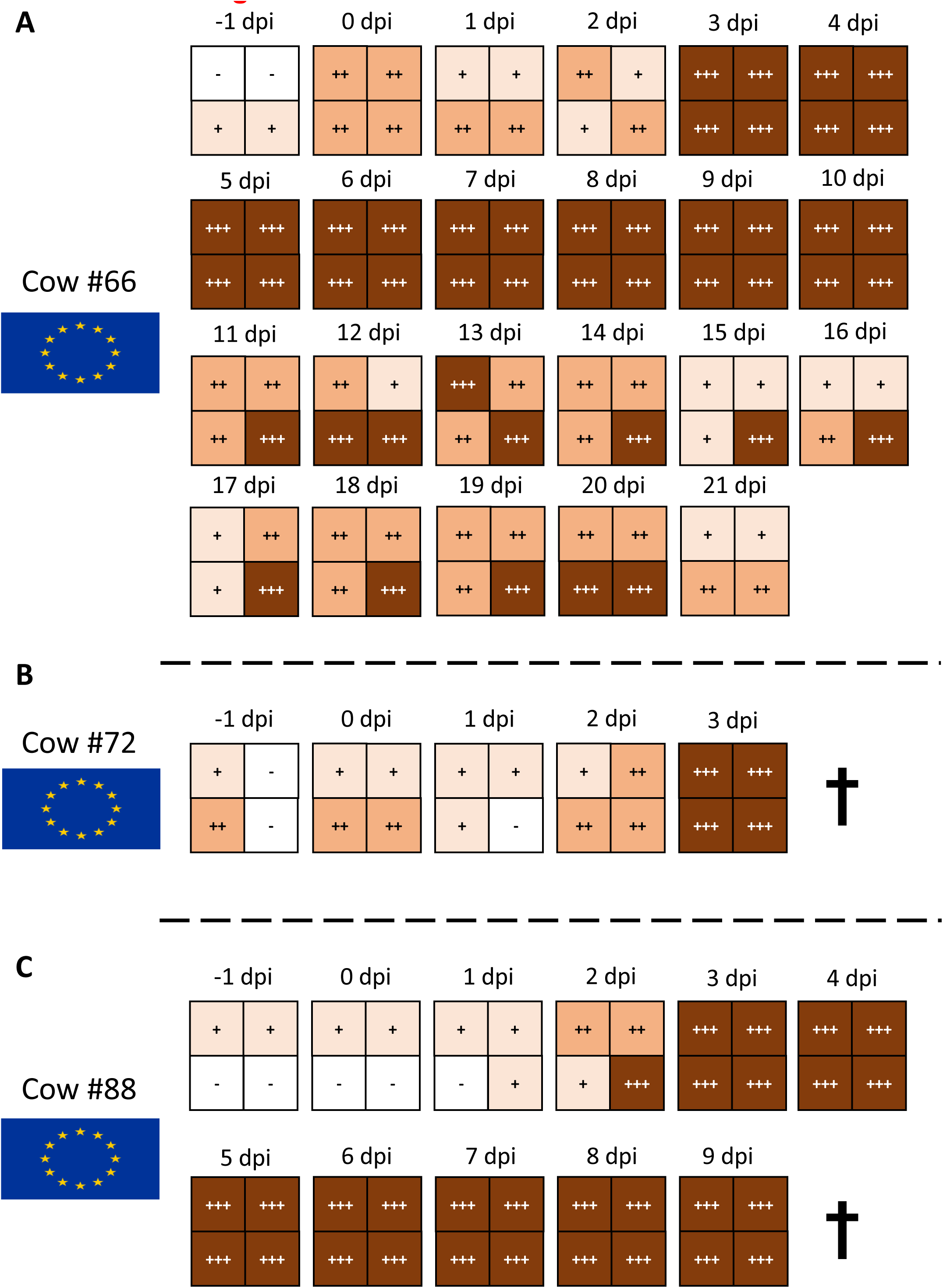
| California Mastitis Test (CMT) of lactating cows infected with H5N1 euDG (EU-group) **A – C** CMT of milk samples from cattle infected with H5N1 euDG during the course of the experiment until their respective euthanasia timepoint.

**Extended Data Fig. 6.**
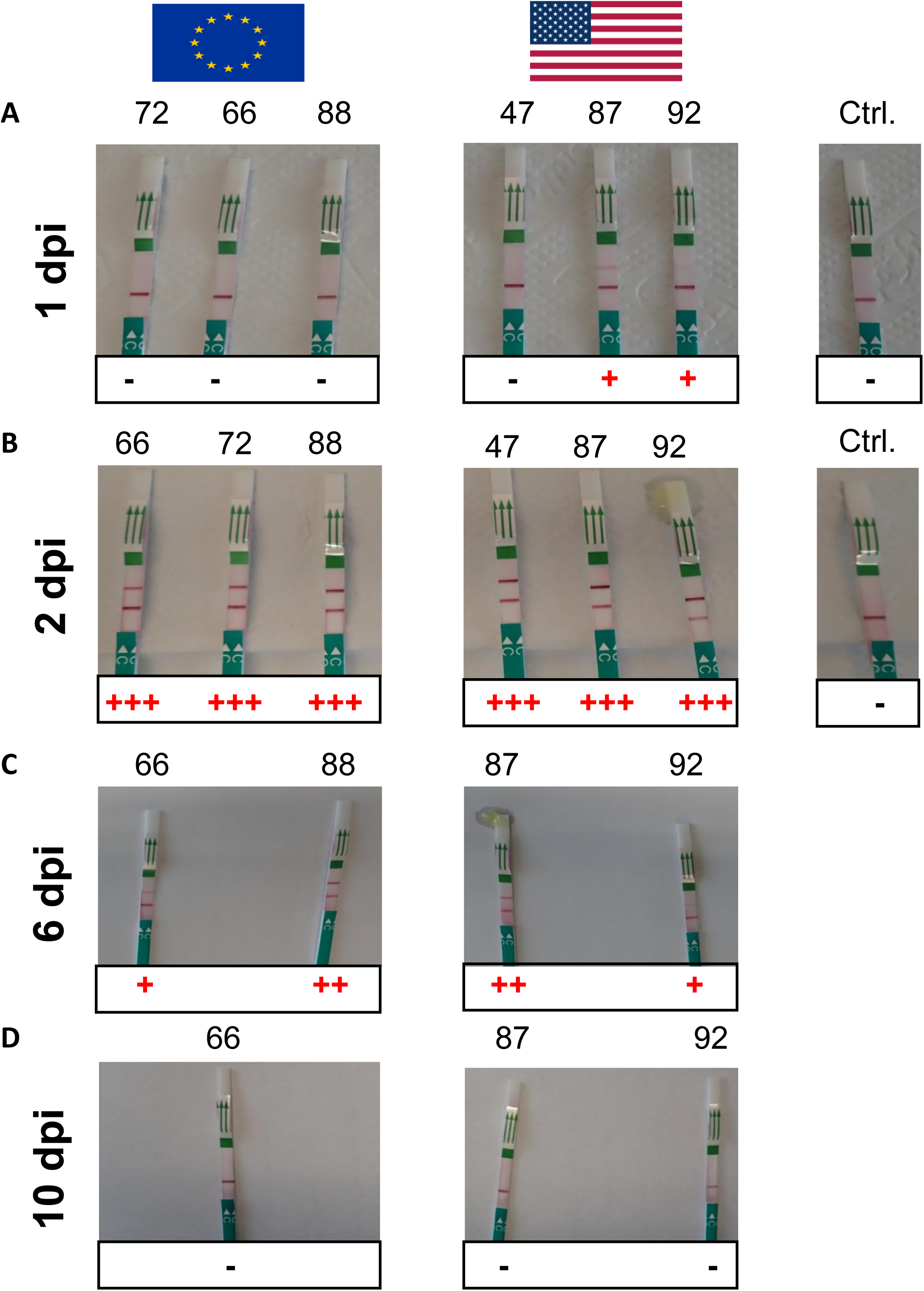
| Megacor-RAT from milk samples of H5N1 experimentally infected dairy cattle. **A** H5-specific Megacor-RAT used for milk samples of H5N1-infected cattle at 1 dpi. Positive samples are depicted with a red cross. **B** H5-specific Megacor-RAT used for milk samples of H5N1-infected cattle at 2 dpi. All H5N1-infected lactating dairy cattle have become positive via the H5-specific RAT from Megacor already at 2 dpi, irrespective of the H5N1-virus isolate used. **C** H5-specific Megacor-RAT used for milk samples of H5N1-infected cattle at 6 dpi. **D** H5-specific Megacor-RAT used for milk samples of H5N1-infected cattle at 10 dpi. All cows have become already negative via the RAT at 10 dpi.

**Extended Data Fig. 7.**
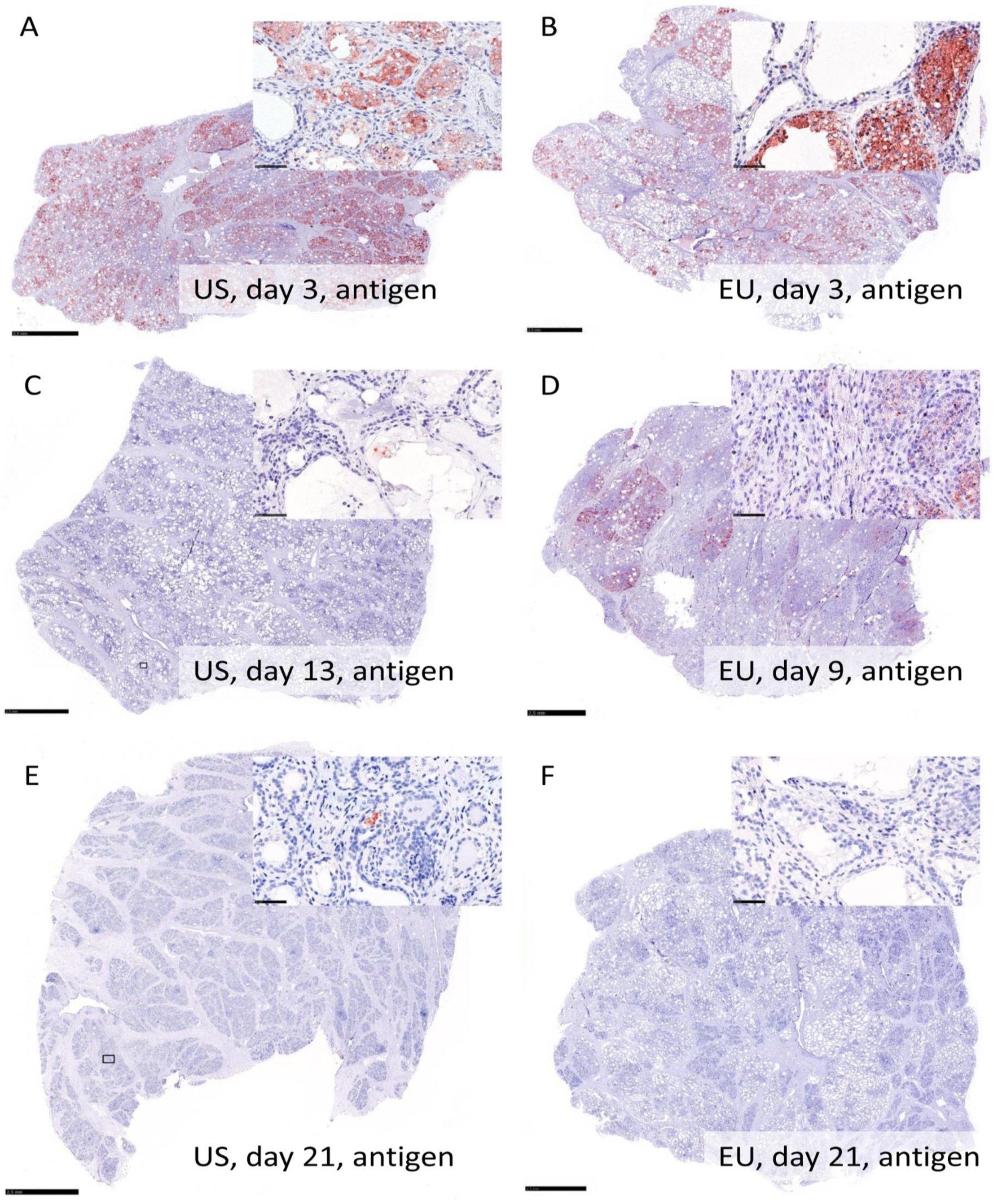
| Influenza A virus nucleoprotein detection using immunohistochemistry in the mammary gland of cattle after intramammary infection with H5N1 B3.13 and H5N1 euDG. The distribution was graded on an ordinal scale with scores 0 = no antigen, 1 = focal, affected cells/tissue <5% or up to 3 foci per tissue; 2 = multifocal, 6%–40% affected; 3 = coalescing, 41%–80% affected; 4 = diffuse, >80% affected. Representative pictures were taken from the most severely affected quarter from each cow. A Score 4, H5N1 B3.13, 3 dpi. B Score 3, H5N1 euDG, 3 dpi. C Score 1, H5N1 B3.13, 13 dpi. D Score 3, H5N1 euDG, 9 dpi. E Score 1, H5N1 B3.13, 21 dpi. F Score 0, H5N1 euDG, 21 dpi. Scale bar 2.5 mm and 50 µm (inlay).

**Extended Data Fig. 8.**
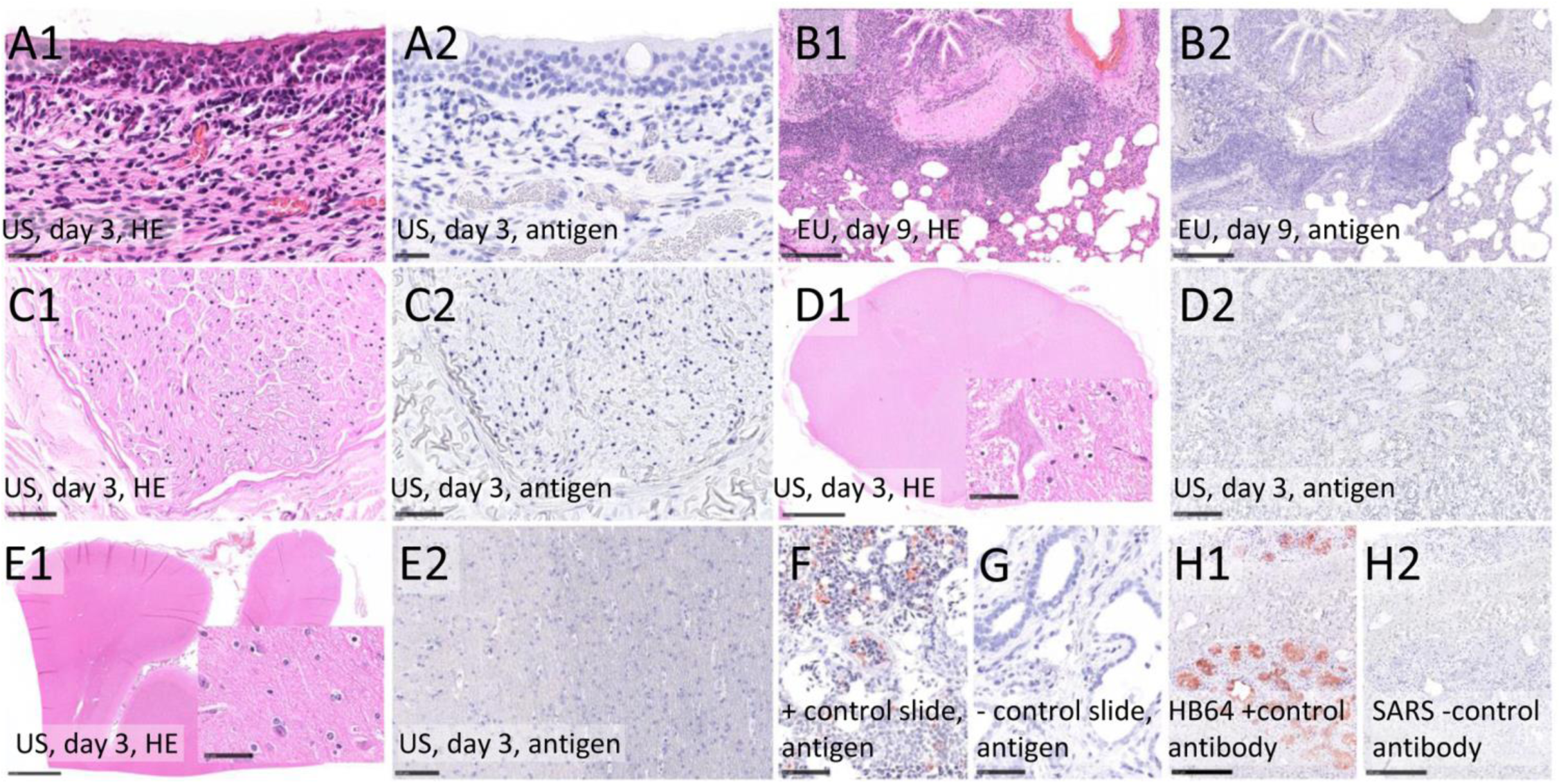
| Histopathology and Influenza A virus nucleoprotein detection (antigen) of cattle after intramammary infection with H5N1 B3.13 and H5N1 euDG including tissue controls. **A** Nasal concha: Chronic-active rhinitis (A1) lacking IAV antigen (A2). **B**: Lung: Chronic bronchointerstitial pneumonia in convalescence phase (B1), lacking IAV NP (B2). **C** Genitofemoral nerve: No findings (C1), no IAV antigen (C2). **D** Spinal cord: No findings (D1), no IAV antigen (D2). **E** Brain, cortex: No findings (E1), no IAV antigen (E2). **F** Positive control slide, HPAIV infected chicken, lung: abundant IAV antigen. **G** Negative control slide, uninfected cow, mammary gland: no IAV antigen. **H1** Mammary gland: cow infected with H5N1 B3.13, 3 dpi, abundant IAV antigen. **H2** Consecutive slide of H1: an irrelevant antibody (anti Sars clone 4F3C4) yielded no immunopositive reaction. Hematoxylin and eosin (HE) stain (**A1, B1, C1, E1**) and immunohistochemistry (antigen) on consecutive (**A2, B2, C2, E2, H1, H2**) or independent (**F, G**) slides. Scale bar 25 µm (**A1-2**), 50 µm (**C1-2**, inlay **D1, E1, F, G**), 100 µm (**D2, E2, H1-2**), 250 µm (**B1-2**), 2.5 mm (**D1, E1**).

**Extended Data Fig. 9.**
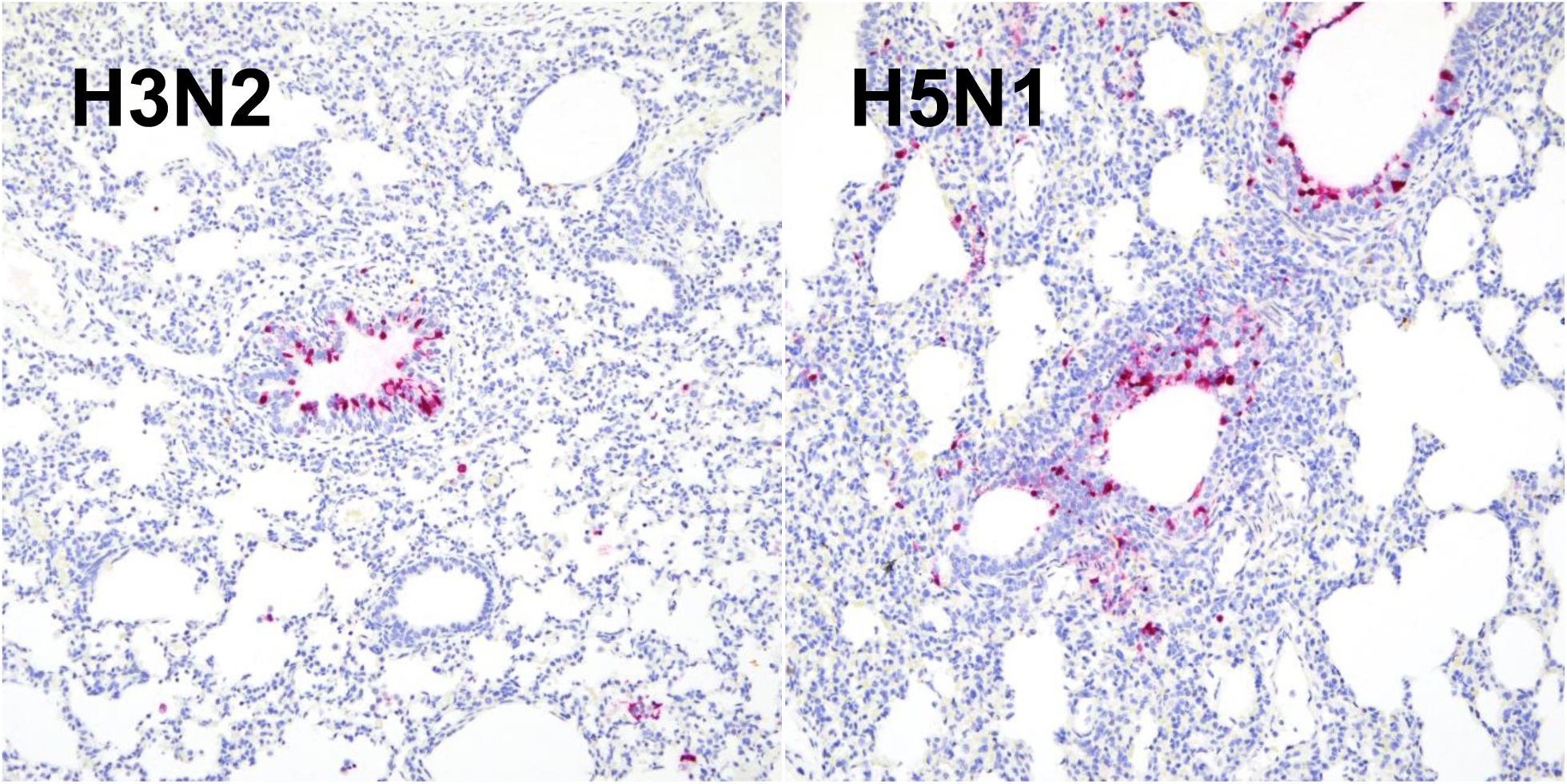
| Tissue controls used for immunohistochemistry. The anti-NP antibody used strongly labels influenza virus A H3N2 and H5N1-infected epithelial cells lining bronchioles.

**Extended Data Fig. 10.**
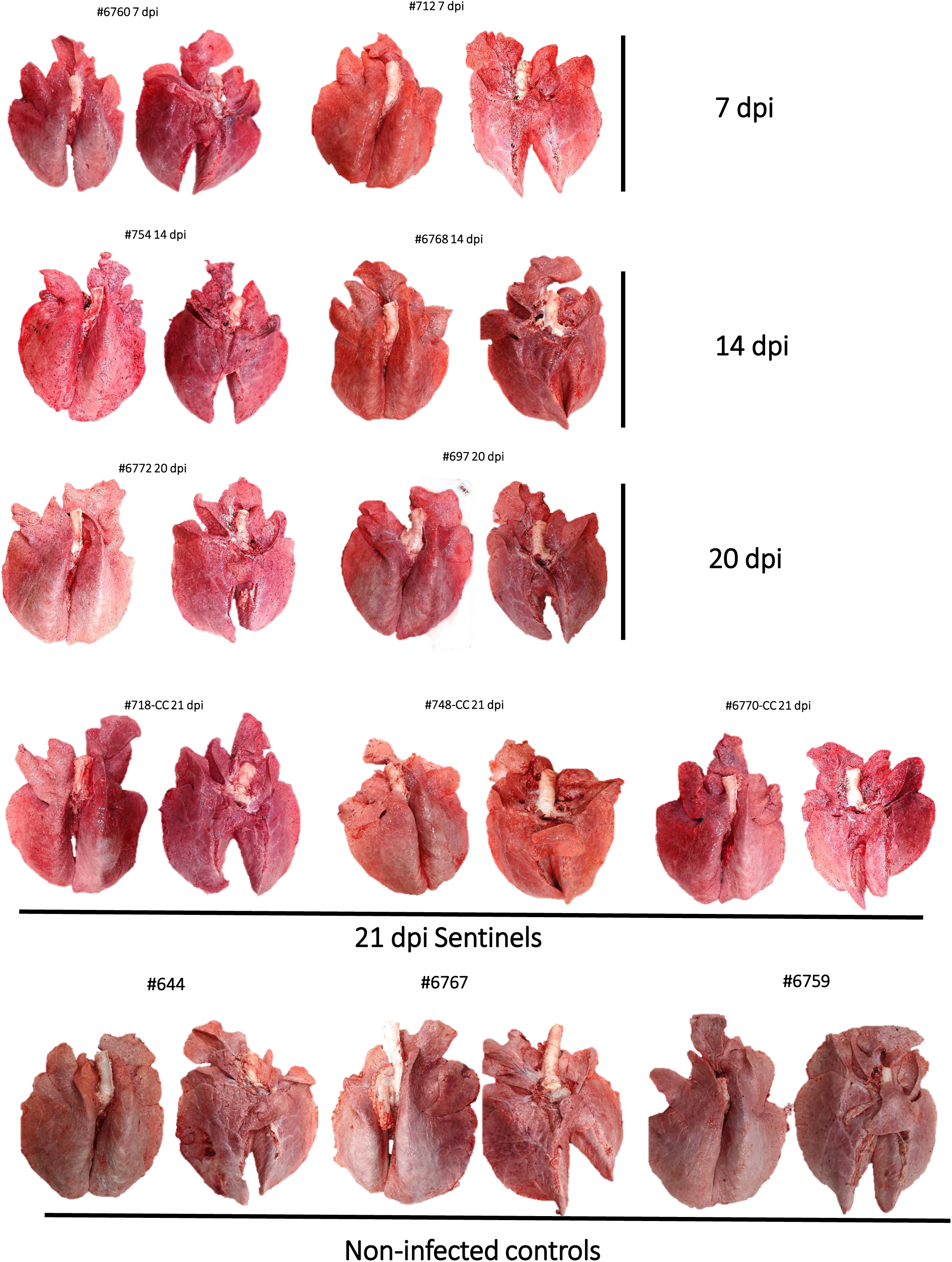
| Gross lung pathology of calves. **(A)** At 7 dpi, multiple well-defined pulmonary lobules were red and slightly depressed on the right cranial lobe (congestion and partial atelectasis) affecting approximately 20% of the cranial and caudal portions of the right cranial lobe extending into the right middle and caudal lobe of one of the two principal-infected calves (#712). There was a focal area of mild subpleural hemorrhage on the ventral surface of the left caudal lung lobe of animal #6760. **(B)** At 14 dpi, one of the two principal-infected calves (#754) had multifocal to coalescing red and depressed foci of congestion and atelectasis on the left and right cranial lobes. Approximately 60% of the caudal portion of the left cranial lobe, 55-60% of both the cranial and caudal portions of the right cranial lobe and <5% of the accessory lobe were affected. There were also multiple pleural adhesions to the thoracic wall. **(C)** At 20 dpi, the two principal-infected calves #6772 and #697) had either few small red and slightly depressed foci of congestion and atelectasis on the left cranial lobes (#6772), or a focal, similar area on the apical portion of the right middle lobe (#697). **(D)** Postmortem examinations of the three sentinel animals were performed at 21 dpi and revealed scattered red foci of pulmonary congestion/atelectasis. In animal #748, there were multiple, small foci of mild consolidation in the left and right cranial lobes (5% of lung affected) and few pleural adhesions to the thoracic cavity. For animal #6770, congestion and atelectasis were accompanied by mild to moderate edema affecting predominately the right lung lobes. **(E)** One of the three negative control calves (#6767) had a small isolated focus of consolidation of the pulmonary parenchyma at the apical margin of the right middle lobe. Gross lesions were not appreciated in the remaining negative control animals.

## Extended Data Tables

**Extended Data Table 1.**
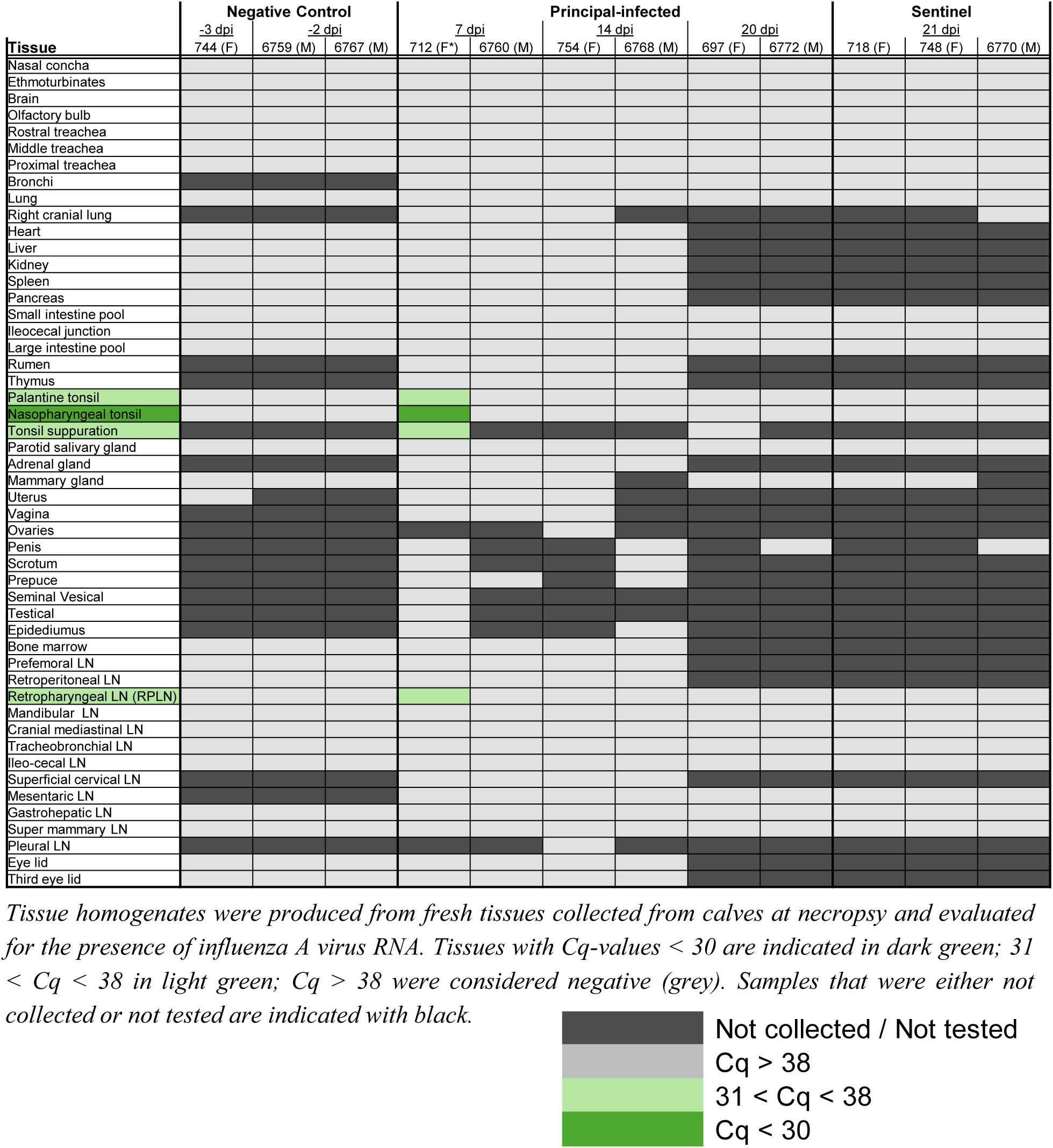
| RT-qPCR results of tissues collected from calves.

**Extended Data Table 2.**
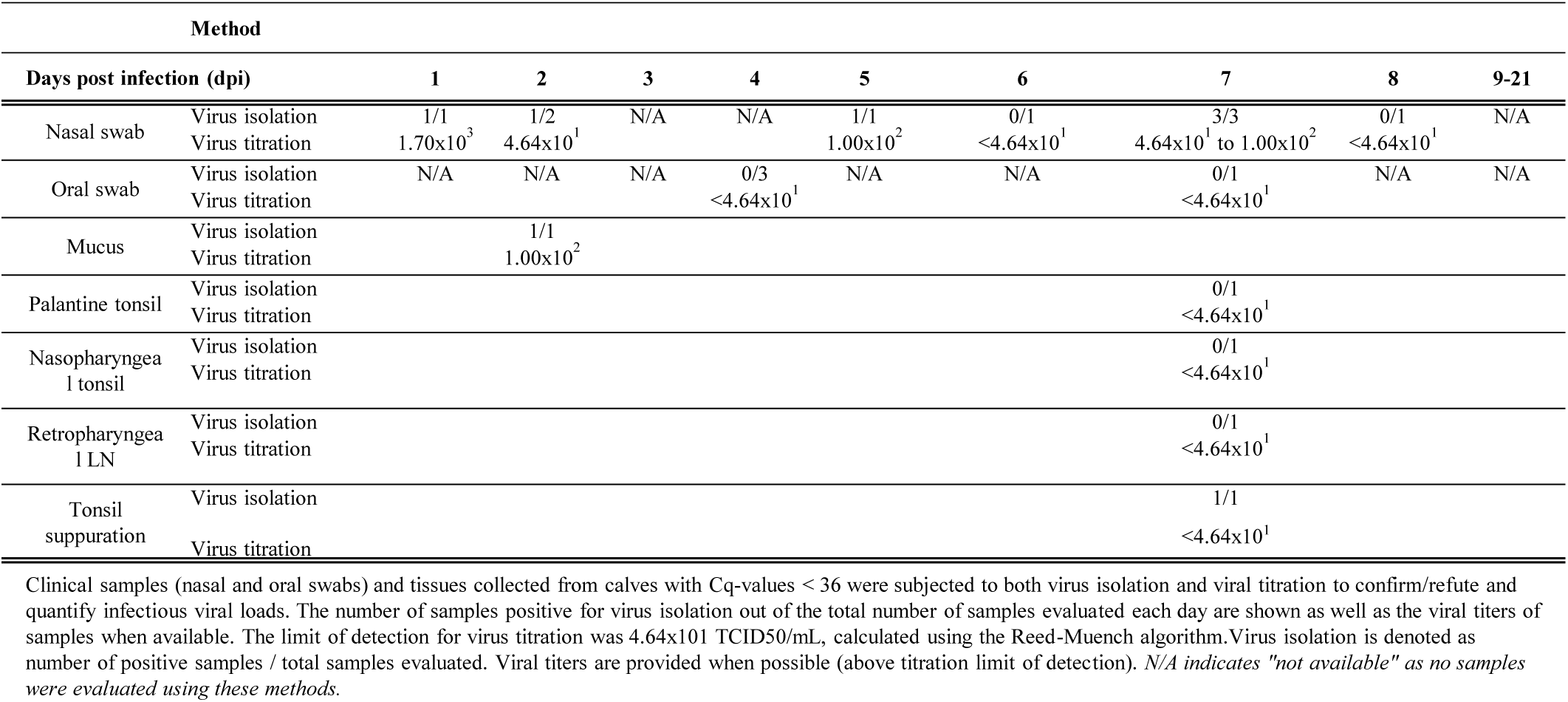
| Viral shedding and isolation in tissues of principal-infected calves following challenge with H5N1 B3.13.

**Extended Data Table 3.** | Results of bovine respiratory disease complex RT-qPCR panel in calves.

|  | ANIMAL ID | Bovine Viral Diarrhea Virus | Bovine Herpesvirus 1 | Bovine Respiratory Syncytial Virus | Mycobacterium bovis | Bovine Corona Virus |
| --- | --- | --- | --- | --- | --- | --- |
| Principal-infected calves | 697 | Negative | Negative | Negative | Negative | Negative |
|  | 712 | Negative | Negative | Negative | Negative | Negative |
|  | 754 | Negative | Negative | Negative | Negative | Negative |
|  | 6760 | Negative | Negative | Negative | Negative | Negative |
|  | 6768 | Negative | Negative | Negative | Negative | Negative |
|  | 6772 | Negative | Negative | Negative | Negative | Negative |
| Sentinel calves | 718 | Negative | Negative | Negative | Negative | Negative |
|  | 748 | Negative | Negative | Negative | Negative | Negative |
|  | 6770 | Negative | Negative | Negative | Negative | Negative |
| Negative control calves | 744 | Negative | Negative | Negative | Negative | Negative |
|  | 6759 | Negative | Negative | Negative | Negative | Negative |
|  | 6767 | Negative | Negative | Negative | Negative | Negative |

|  | ANIMAL ID | Influenza D Virus | Mannheimia haemolytica | Pasteurella multocida | Histophilus somni | Bibersteinia trehalosi |
| --- | --- | --- | --- | --- | --- | --- |
| Principal-infected calves | 697 | Negative | Negative | 38.4 | Negative | Negative |
|  | 712 | Negative | Negative | 31.26 | Negative | 36.25 |
|  | 754 | Negative | 33.07 | 36.18 | Negative | Negative |
|  | 6760 | Negative | Negative | Negative | Negative | Negative |
|  | 6768 | 32.96 | 32.67 | 34.54 | Negative | Negative |
|  | 6772 | Negative | Negative | 36.46 | Negative | 33.49 |
| Sentinel calves | 718 | Negative | Negative | Negative | Negative | 37.27 |
|  | 748 | Negative | 28.77 | 34.29 | Negative | 35.64 |
|  | 6770 | 38.09 | Negative | Negative | Negative | Negative |
| Negative control calves | 744 | Negative | Negative | 35.78 | Negative | 35.58 |
|  | 6759 | 36.37 | Negative | 36.43 | Negative | 38.46 |
|  | 6767 | 19.7 | 31.44 | 31.69 | Negative | 34.4 |

Nasal swabs collected from calves at -8 dpi were submitted to the Kansas State Veterinary Diagnostic Laboratory for comprehensive screening of common bovine respiratory disease complex (BRDC) pathogens using qPCR/RT-qPCR detection methods. Interpretation of results: Positive = Ct values <36; Suspect/Inconclusive = Ct values between 36 and 39; Negative = Ct values > 39 or 0. Clinical samples (nasal and oral swabs ) and RNA from a limited set of post mortem samples were tested for the present of Influenza D PCR as a single-plex assay and were negative<sup>44</sup> .

**Extended Data Table 4.** | Gross lung scores for calves.

|  | Calf ID | Gender | dpi | Percentage of individual lung lobes affected |  |  |  |  |  |  | Average percentage |  |  |
| --- | --- | --- | --- | --- | --- | --- | --- | --- | --- | --- | --- | --- | --- |
|  |  |  |  | LT CR | LT M | LT CD | RT CR | RT M | RT CD | A | Left | Right | Total |
| Principal-infected calves | 6760 | Male | 7 | 5 | 5 | 5 | 5 | 5 | 10 | 5 | 5 | 6 | 6 |
|  | 712 | Female* | 7 | 5 | 15 | 10 | 30 | 20 | 10 | 20 | 10 | 20 | 16 |
|  | 6768 | Male | 14 | 5 | 5 | 0 | 15 | 10 | 5 | 10 | 3 | 10 | 7 |
|  | 754 | Female | 14 | 4 | 60 | 5 | 55 | 60 | 5 | 25 | 23 | 36 | 31 |
|  | 6772 | Male | 20 | 5 | 20 | 0 | 0 | 5 | 0 | 0 | 8 | 1 | 4 |
|  | 697 | Female | 20 | 0 | 5 | 5 | 0 | 0 | 0 | 0 | 3 | 0 | 1 |
| Sentinel calves | 718 | Female | 21 | 0 | 0 | 0 | 0 | 0 | 0 | 0 | 0 | 0 | 0 |
|  | 748 | Female | 21 | 5 | 5 | 0 | 5 | 0 | 0 | 0 | 3 | 1 | 2 |
|  | 6770 | Male | 21 | 20 | 30 | 10 | 10 | 10 | 5 | 5 | 20 | 8 | 13 |
| Negative control calves | 744 | Female | -2 | 5 | 0 | 0 | 5 | 0 | 0 | 0 | 2 | 1 | 1 |
|  | 6759 | Male | -3 | 0 | 0 | 0 | 0 | 0 | 0 | 0 | 0 | 0 | 0 |
|  | 6767 | Male | -3 | 0 | 0 | 0 | 5 | 10 | 0 | 0 | 0 | 4 | 2 |

Gross lung scores: Percent of lung affected with gross lesions including congestion with atelectasis or edema, pneumonia, hemorrhage, and plural fibrosis when present. Evaluation of individual lung lobes from calves, reported as the percentage of lung affected with gross lesions including congestion with atelectasis or edema, pneumonia, hemorrhage, and plural fibrosis when present. Total percentage of lung affected is listed in the final column. LT CR= left cranial lung lobe; LT M=left middle lung lobe; LT CD= left caudal lung lobe; RT CR= right cranial lung lobe; RT M= right middle lung lobe; RT CD= right caudal lung lobe; A= accessory lobe \* indicates one calf as hermaphroditic.

**Extended Data Table 5.** | Tissue samples from intramammary infected cows and methods applied including hematoxylin-eosin stain (HE) and immunohistochemical Influenza virus nucleoprotein detection (IHC)

| Animal ID# | 80 | 72 | 47 | 88 | 92 | 87 | 66 |
| --- | --- | --- | --- | --- | --- | --- | --- |
| Virus | mock | EU | US | EU | US | US | EU |
| Autopsy dpi | 3 | 3 | 3 | 9 | 13 | 21 | 21 |
| Tissues |  |  |  |  |  |  |  |
| Mammary quarter, left, front | HE&IHC | HE&IHC | HE&IHC | HE&IHC | HE&IHC | HE&IHC | HE&IHC |
| Mammary quarter, right, front | HE&IHC | HE&IHC | HE&IHC | HE&IHC | HE&IHC | HE&IHC | HE&IHC |
| Mammary quarter, left, rear | HE&IHC | HE&IHC | HE&IHC | HE&IHC | HE&IHC | HE&IHC | HE&IHC |
| Mammary quarter, right, rear | HE&IHC | HE&IHC | HE&IHC | HE&IHC | HE&IHC | HE&IHC | HE&IHC |
| Teat, left, front | HE | HE&IHC | HE&IHC | HE&IHC | HE&IHC | HE&IHC | HE&IHC |
| Teat, right, front | HE | HE&IHC | HE&IHC | HE&IHC | HE&IHC | HE&IHC | HE&IHC |
| Teat, left, rear | HE | HE&IHC | HE&IHC | HE&IHC | HE&IHC | HE&IHC | HE&IHC |
| Teat, right, rear | HE | HE&IHC | HE&IHC | HE&IHC | HE&IHC | HE&IHC | HE&IHC |
| Nose, vestibulum (squamous mucosa) | HE | HE&IHC | HE&IHC | HE&IHC | HE&IHC | - | - |
| Nasal conchae, respiratory | HE | HE&IHC | HE&IHC | HE&IHC | HE&IHC | - | - |
| Nasal conchae, olfactory | HE | HE&IHC | HE&IHC | HE&IHC | HE&IHC | - | - |
| Trachea | HE | HE&IHC | HE&IHC | HE&IHC | HE | - | - |
| Lung, left, cranial lobe (cranial part) | HE | HE&IHC | HE&IHC | HE&IHC | HE | - | - |
| Lung, left, cranial lobe (caudal part) | HE | HE&IHC | HE&IHC | HE&IHC | HE | - | - |
| Lung, left, caudal lobe | HE | HE&IHC | HE&IHC | HE&IHC | HE | - | - |
| Lung, right, cranial lobe (cranial part) | HE | HE&IHC | HE&IHC | HE&IHC | HE | - | - |
| Lung, right, cranial lobe (caudal part) | HE | HE&IHC | HE&IHC | HE&IHC | HE | - | - |
| Lung, right, mid lobe | HE | HE&IHC | HE&IHC | HE&IHC | HE | - | - |
| Lung, right, caudal lobe | HE | HE&IHC | HE&IHC | HE&IHC | HE | - | - |
| Lung, accessory lobe | HE | HE&IHC | HE&IHC | HE&IHC | HE | - | - |
| Nerve, genitofemoralis | HE | HE&IHC | HE&IHC | HE&IHC | HE&IHC | - | - |
| Brain, olfactory bulb | HE | HE&IHC | HE&IHC | HE&IHC | HE&IHC | - | - |
| Brain, cortex (level of hippocampus) | HE | HE&IHC | HE&IHC | HE&IHC | HE&IHC | - | - |
| Brain, cerebellum | HE | HE&IHC | HE&IHC | HE&IHC | HE&IHC | - | - |
| Brain, brainstem, medulla oblongata | HE | HE&IHC | HE&IHC | HE&IHC | HE&IHC | - | - |
| Spinal cord (cervical) | HE | HE&IHC | HE&IHC | HE&IHC | HE&IHC | - | - |
| Spinal cord (thoracic) | HE | HE&IHC | HE&IHC | HE&IHC | HE&IHC | - | - |
| Spinal cord (lumbar) | HE | HE&IHC | HE&IHC | HE&IHC | HE&IHC | - | - |
| Heart | HE | HE&IHC | HE&IHC | HE&IHC | HE | - | - |
| Liver | HE | HE&IHC | HE&IHC | HE&IHC | HE | - | - |
| Spleen | HE | HE&IHC | HE&IHC | HE&IHC | HE | - | - |
| Lymph node, tracheobronchial | HE | HE&IHC | HE&IHC | HE&IHC | HE | - | - |
| Lymph node, supramammary | HE | HE&IHC | HE&IHC | HE&IHC | HE | - | - |
| Tonsil, pharyngeal | HE | HE&IHC | HE&IHC | HE&IHC | HE | - | - |
| Tonsil, palatine | HE | HE&IHC | HE&IHC | HE&IHC | HE | - | - |
| Kidney | HE | HE&IHC | HE&IHC | HE&IHC | HE | - | - |
| Urinary bladder | HE | HE&IHC | HE&IHC | HE&IHC | HE&IHC | - | - |
| Uterus | HE | HE&IHC | HE&IHC | HE&IHC | HE | - | - |
| Cervix | HE | HE&IHC | HE&IHC | HE&IHC | HE | - | - |
| Vagina (vestibulum) | HE | HE&IHC | HE&IHC | HE&IHC | HE | - | - |
| Skin, inguinal | HE | HE&IHC | HE&IHC | HE&IHC | HE | - | - |
| Duodenum | HE | HE&IHC | HE&IHC | HE&IHC | HE | - | - |
| Rektum | HE | HE&IHC | HE&IHC | HE&IHC | HE | - | - |
| Pancreas | HE | HE&IHC | HE&IHC | HE&IHC | HE | - | - |
| Adrenal gland | HE | HE&IHC | HE&IHC | HE&IHC | HE | - | - |

## Supplementary Data 1: Lactating dairy cows, summary of relevant findings in tissues; interpreted to be not associated with IAV-infection

#80, negative control animal

- Thorax: pleural adhesion, right caudal lung lobe
- Lung, right, caudal lobe: pleuritis, chronic, focal, moderate, lymphoplasmacytic and fibrosing; pneumonia, bronchointerstitial, chronic-active, multifocal to coalescing, moderate, mainly interstitial, lymphoplasmacytic with prominent hyperplasia/hypertrophy of bronchial epithelium and type II pneumocytes, some areas showing suppurative-necrotizing bronchitis and bronchiolitis, with intraluminal cellular debris, mucin, proteinaceous edema, bronchus-associated lymphoid tissue (BALT) hyperplasia
- Spleen: hyperemia, acute, diffuse, severe, moderate number of hemosiderin-laden macrophages (intracytoplasmic, pale brown pigment, consistent with hemosiderin); minimally increased number of neutrophils within sinuses
- Heart: atrial appendages, epicarditis and pericarditis, chronic, mild, focal, fibrosing; myocarditis, chronic-active, focal, mild, lymphoplasmacytic and eosinophilic
- Liver: abscess, focal, up to 15 cm in diameter; amyloidosis, periportal and bridging, mild
- Kidney: nephritis, chronic, interstitial, multifocal, mild, lymphoplasmacytic and histiocytic
- Small intestine: lamina propria mucosae with high numbers of lymphoplasmacytic infiltrates, fewer macrophages and scattered eosinophils, mucosal epithelium intact
- Spinal cord: perivascular cuffing, lymphocytic, focal, minimal
- Brain: perivascular macrophages, oligofocal, minimal, with intracytoplasmic, pale brown pigment (consistent with hemosiderin)
- No pathogens, or inclusion bodies, or syncytia found
- Other tissues: no relevant findings, immunohistochemistry for influenza A virus nucleoprotein (IAV NP): all negative

#47, US isolate, 3 dpi

- Nasal conchae mucosa/pharynx: petechia, multifocal, mild; rhinitis, chronic-active, diffuse, moderate, mainly lymphoplasmacytic, some areas with neutrophilic infiltrates, many mucosal transmigrating neutrophils; ciliated respiratory epithelium mostly intact, some areas with loss of cilia and/or degeneration and single cell necrosis
- Tonsil, pharyngeal / palatine: tonsillitis, acute, diffuse, moderate, suppurative and hemorrhagic, with prominent intraluminal debris, admixed with foreign material, intralesional bacteria
- Trachea: tracheitis, acute, diffuse, mild, necrotizing and suppurative, with luminal cellular debris and proteinaceous material
- Spleen: hyperemia, acute, diffuse, severe, moderate number of hemosiderin-laden macrophages
- Lymph node, tracheobronchial: lymphadenitis, acute, diffuse, mild, with increased number of neutrophils in sinuses
- Heart: atrial appendages, epicarditis and pericarditis, chronic, mild, focal, fibrosing
- Liver: hepatitis, chronic-active, focal, minimal, granulomatous and eosinophilic; increased number of neutrophils in hepatic sinuses and blood vessels
- Kidney: nephritis, chronic, interstitial, multifocal, mild, lymphoplasmacytic and histiocytic
- Brain: perivascular macrophages, oligofocal, minimal, with intracytoplasmic, pale brown pigment (consistent with hemosiderin)
- Brain, olfactory bulb: perivascular glial cell aggregation, focal, minimal
- Adrenal: adrenalitis, chronic, multifocal, mild, lymphocytic
- No further pathogens, or inclusion bodies, or syncytia found
- Other tissues: no relevant findings, immunohistochemistry for influenza A virus nucleoprotein (IAV NP): all negative

#92, US-isolate, 13 dpi

- Nasal conchae: rhinitis, acute, diffuse, mild, suppurative
- Lung, left and right caudal lobe, right cranial lobe (cranial part), accessory lobe: pleuritis, chronic, focal, moderate, lymphoplasmacytic and fibrosing
- Spleen: hyperemia, acute, diffuse, severe, moderate number of hemosiderin-laden macrophages; minimally increased number of neutrophils within sinuses; follicular hyperplasia, mild
- Lymph node, tracheobronchial and iliac: follicular hyperplasia, mild
- Liver: perihepatitis, chronic, focal, mild, fibrosing
- Kidney: nephritis, chronic, interstitial, moderate, lymphoplasmacytic, some areas with prominent fibrosis and glomerulosclerosis and/or tubular degeneration and regeneration
- Small intestine: lamina propria mucosae with high numbers of lymphoplasmacytic infiltrates, fewer macrophages and scattered eosinophils, mucosal epithelium intact
- Brain: perivascular macrophages, oligofocal, minimal, with intracytoplasmic, pale brown pigment (consistent with hemosiderin)
- No pathogens, or inclusion bodies, or syncytia found
- Other tissues: no relevant findings, immunohistochemistry for influenza A virus nucleoprotein (IAV NP): all negative

#87, US-isolate, 21 dpi

- Liver: perihepatitis, chronic, focal, mild, fibrosing
- Histopathology done for mammary gland and teat only; immunohistochemistry for influenza A virus nucleoprotein (IAV NP): negative

#72, EU-Isolate, 3 dpi

- Thorax: Pleural adhesion, left and right cranial lung lobe
- Lung left and right cranial lobes, left caudal lobe, accessory lobe: pleuritis, chronic, focal, moderate to severe, lymphoplasmacytic and fibrosing; pneumonia, bronchointerstitial, chronic-active, multifocal to coalescing, moderate to severe, mainly interstitial, lymphoplasmacytic with prominent hyperplasia/hypertrophy of bronchial epithelium and type II pneumocytes, some areas showing suppurative-necrotizing bronchitis and bronchiolitis, with intraluminal cellular debris, mucin, proteinaceous edema, bronchus-associated lymphoid tissue (BALT) hyperplasia
- Lymph node, tracheobronchial: lymphadenitis, acute, diffuse, moderate, with increased number of neutrophils in sinuses, with follicular hyperplasia, mild
- Spleen: hyperemia, acute, diffuse, severe, moderate number of hemosiderin-laden macrophages; minimally increased number of neutrophils within sinuses;
- Liver: perihepatitis, chronic, focal, mild, fibrosing; amyloidosis, periportal and bridging, moderate; minimally increased number of neutrophils in hepatic sinuses and blood vessels; single cell necrosis/apoptosis, hepatocellular, multifocal, mild
- Kidney: amyloidosis, interstitial, moderate
- Adrenal gland: amyloidosis, moderate
- Small and large intestine: lamina propria mucosae with high numbers of lymphoplasmacytic infiltrates, fewer macrophages and scattered eosinophils, mucosal epithelium intact
- Brain: perivascular macrophages, oligofocal, minimal, with intracytoplasmic, pale brown pigment (consistent with hemosiderin)
- Brain stem: perivascular cuffing, lymphocytic, focal, minimal
- No pathogens, or inclusion bodies, or syncytia found
- Other tissues: no relevant findings, immunohistochemistry for influenza A virus nucleoprotein (IAV NP): all negative

#88, EU-isolate, 9 dpi

- Thorax: Pleural adhesion, left and right cranial lung lobes
- Lung, all lung lobes: pneumonia, interstitial, chronic, diffuse, moderate, lymphoplasmacytic and histiocytic, with moderate BALT hyperplasia and perivascular lymphocytic hyperplasia, moderate interstitial/pleural fibrosis, some areas with moderate II pneumocytes hyperplasia
- Spleen: hyperemia, acute, diffuse, severe, moderate number of hemosiderin-laden macrophages; minimally increased number of neutrophils within sinuses
- Kidney: fibrosis, chronic, interstitial, minimal
- Uterus: endometritis, subacute, diffuse, moderate, suppurative, mucosal epithelium intact
- Rumen and omasum: Erosions, acute, multifocal, mild
- Brain, cortex: perivascular glial cell aggregation, focal, minimal
- Brain: perivascular macrophages, oligofocal, minimal, with intracytoplasmic, pale brown pigment (consistent with hemosiderin)
- Small intestine: lamina propria mucosae with high numbers of lymphoplasmacytic infiltrates, fewer macrophages and scattered eosinophils, mucosal epithelium intact
- Large intestine: Proctitis, subacute, multifocal, severe, ulcerative, with early granulation tissue formation and focal vascular fibrinoid necrosis
- No pathogens, or inclusion bodies, or syncytia found
- Other tissues: no relevant findings, immunohistochemistry for influenza A virus nucleoprotein (IAV NP): all negative

#66, EU-isolate, 21 dpi

- Thorax: Pleural adhesion, left and right cranial lung lobes
- Heart: atrial appendages, epicarditis and pericarditis, chronic, mild, focal, fibrosing;
- Histopathology done for mammary gland and teat only; immunohistochemistry for influenza A virus nucleoprotein (IAV NP): negative

